# Investigating the Dynamic Relationship Between Anxiety and Spatial Memory Using Autonomous Ecological Momentary Assessment

**DOI:** 10.64898/2026.05.15.725563

**Authors:** Claire Z. Han, Kun C. Zhao, Linda M. Wang, Hongkun Zhu, Yangjia Li, Luca D. Kolibius, Angela G. Velazquez, Sandy Song, Amy Cami, Jerina Carmona, Marla Hamberger, Randy P. Auerbach, Catherine A. Schevon, Joshua Jacobs, Brett E. Youngerman

## Abstract

Anxiety has been extensively studied in relation to memory, yet its dynamic association with spatial episodic memory in naturalistic clinical settings remains largely unexplored. We developed an anxiety-spatial-memory EMA protocol (asm-EMA) and deployed it in 30 epilepsy patients undergoing inpatient EEG monitoring, delivering combined momentary anxiety ratings and a validated spatial memory task pseudo-randomly every 90–150 minutes across multiple days. Subject-level asm-EMA average and session-to-session variability correlated with standard neuropsychological assessments, supporting the clinical validity of the protocol. Momentary anxiety was not associated with reduced spatial memory accuracy. Higher within-person STAI-6 was associated with faster retrieval responses, but this association was attenuated after accounting for prior task exposure, suggesting that response-time effects may partly reflect task familiarity. Temporal analyses identified both gradual hospital-day change and short-term carry-over across anxiety, pain, and spatial memory measures. STAI-6 anxiety and spatial memory showed stronger hospital-day trends, whereas VAS anxiety showed the clearest carry-over from one session to the next. The asm-EMA protocol provides a feasible, autonomous framework for capturing moment-to-moment anxiety-memory dynamics in naturalistic settings.

## Introduction

It is well established that anxiety profoundly influences cognitive function, most notably memory. Indeed, attentional control theory^1^ posits that elevated anxiety competes for the limited attentional resources required for successful memory encoding and retrieval. Meta-analytic work demonstrates a reliable negative association between self-reported measures of anxiety and working memory capacity^2^. Further, anxiety disorders are associated with episodic memory impairment^3^, and anxiety symptoms prospectively predict memory decline in cognitively healthy older adults^4^. Spatial working memory appears particularly vulnerable to anxiety^5,6^, consistent with the hippocampus serving a dual role in both spatial cognition and anxiety regulation^7^. Yet despite this body of evidence, the anxiety-memory relationship is far from settled. The working memory impairment robustly observed under controlled laboratory conditions does not replicate when anxiety is measured naturalistically in community samples^8^, suggesting that documented effects may be driven primarily by clinical-level or experimentally induced anxiety rather than the everyday fluctuations most people actually experience. The effect is also paradigm-specific, with anxious individuals showing a recall bias for threatening material but no reliable bias in implicit memory or recognition^9^. Procedural factors such as retention interval further modulate any effect that does emerge, such that higher anxious arousal impairs memory across a waking retention interval but not when retention spans nocturnal sleep^10^. Together, these findings suggest that anxiety does not uniformly impair memory.

This heterogeneity may reflect a structural limitation in how the anxiety-memory relationship has been studied. The bulk of existing evidence derives from single-session, between-subjects designs in which anxiety is either experimentally induced or indexed once as a trait, and memory is probed under tightly controlled conditions far removed from the sustained, fluctuating affective contexts of daily life. Such designs cannot resolve how momentary anxiety co-varies with memory performance within individuals as it fluctuates across hours and days. Whether the between-person associations measured in the laboratory index the same processes as within-person fluctuations in naturalistic settings remains an open question, and the two levels of analysis are known to diverge substantially in magnitude, and at times in direction^11,12^.

Characterizing the within-person relationship between anxiety and memory requires not only repeated measurement but is best studied with a within-person structure, in which each subject serves as their own control and the effects of momentary anxiety are partitioned from stable individual differences such as trait anxiety or baseline cognitive ability^13^. This matters because whether a person’s memory is worse on their more anxious days is a fundamentally different question from whether more anxious people perform worse overall, and collapsing these levels of analysis yields misleading estimates of effect size and generalizability^14^. Ecological momentary assessment (EMA) characterizes a subject’s state at a particular moment in time in a natural environment^15^, and evidence from naturalistic studies confirms it detects effects that laboratory paradigms cannot. In one two-week smartphone EMA study, morning stress anticipation predicted working memory deficits later that day over and above stressful events that actually occurred, a finding that no single-session design could have produced^16^. EMA is therefore well suited to characterizing how anxiety and memory co-vary within individuals across time.

Spatial episodic memory is a particularly informative target because it depends on hippocampal systems that support spatial-contextual binding^17^ and are sensitive to stress-related neuroendocrine and arousal processes relevant to anxiety-memory associations^18,19,20^. Consistent with this framework, animal studies have shown that chronic stress can impair spatial memory performance^21^. Although chronic stress in animal models and momentary anxiety in patients are related but distinct constructs, these findings motivate investigating whether spatial episodic memory performance covaries with anxiety within individuals.

Prior studies have demonstrated the feasibility of repeated cognitive assessment in daily life by incorporating measures of working memory, visuospatial attention, and processing speed into EMA protocols^22, 23^. Extending this approach to spatial episodic memory requires preserving the demands of object-location encoding and retrieval within brief tasks suitable for repeated, self-administered assessment. In our view, the challenge of adapting spatial episodic memory tasks to EMA lies in preserving the demands of object-location encoding and retrieval across repeated assessments while minimizing practice effects, maintaining comparable task difficulty, and accommodating the perceptual and motor constraints of mobile testing.

The inpatient Epilepsy Monitoring Unit (EMU) provides an opportunity to study anxiety-memory relationships during hospitalization alongside continuous clinical video and EEG monitoring. Qualitative research has identified restricted activity, boredom, and waiting for seizures as sources of discomfort during EMU admission^24^. These observations motivated our investigation of whether anxiety varies within individuals during hospitalization and whether such variation is associated with spatial episodic memory performance. Repeated assessment in this setting must also accommodate clinical care without overburdening patients. Prior work supports the feasibility of intensive mobile assessment in clinical populations. Mobile mood assessment has been validated against standard clinical measures in outpatient settings^25,26^, and repeated psychological self-assessment has been implemented successfully in an EMU, albeit without an accompanying cognitive measure^27^. What remains unclear is whether repeated affective sampling can be paired with performance-based spatial memory assessment to test anxiety-memory associations within individuals under naturalistic inpatient conditions.

The present study addresses this gap by combining intensive within-person EMA with a performance-based spatial episodic memory task in patients undergoing multi-day inpatient EMU monitoring, a setting in which sustained, naturalistic clinical stress unfolds in real time. Here we report findings from an asm-EMA (anxiety-spatial-memory EMA) protocol in which 30 EMU inpatients completed repeated assessments across multiple days of hospitalization. Each assessment combined momentary anxiety ratings (VAS and STAI-6) with a validated spatial episodic memory task^28,29^. As an observational study, the present work characterizes how anxiety and spatial memory co-vary under naturalistic clinical stress and establishes asm-EMA protocol as a valid longitudinal affective-cognitive assessment tool in this population. We address three related questions. First, whether the asm-EMA protocol captures clinically meaningful temporal variation in anxiety and spatial memory, as evaluated against standard neuropsychological assessments. Second, whether momentary anxiety is associated with concurrent spatial episodic memory performance, including both between-person differences and within-person fluctuations. Third, whether repeated asm EMA measurements capture short term temporal dynamics, including persistence of anxiety and memory states across adjacent sessions and lagged associations between anxiety and subsequent memory performance.

The asm-EMA protocol is a core component of the broader Context-Aware Multimodal Ecological Research and Assessment platform, which integrates asm-EMA behavioral data with synchronized continuous intracranial recordings, physiological monitoring, and passively acquired data streams including audio, video, and smartphone digital phenotyping^30^. The behavioral signals from asm-EMA sessions anchor these multimodal data streams, enabling the platform to identify the neural and physiological correlates of anxiety-memory states in real time and ultimately to trigger closed-loop neuromodulation of anxiety at behaviorally informed moments. Establishing that asm-EMA protocol reliably captures meaningful within-person variation in anxiety and spatial memory is therefore a necessary first step toward that larger goal.

## Results

### asm-EMA Protocol Feasibility and Compliance

Over the course of hospitalization, 496 asm-EMA sessions were delivered across 30 inpatients (described in Table 1). Of these, 424 sessions, or 85.5%, were at least partially completed. Partially or fully completed sessions were distributed across three completion tiers. Sixty-eight sessions, or 13.7% of all prompts, included VAS ratings only. Eighty-five sessions, or 17.1% of all prompts, included the full anxiety assessment with VAS and STAI-6 but did not proceed to the spatial memory task. The remaining 271 sessions, or 54.6% of all prompts, included the full protocol with VAS, STAI-6, and the Treasure Hunt spatial memory task. Seven full-protocol sessions were excluded because of inattentive responding to the self-reported anxiety items, yielding 264 complete sessions for analysis. 28 participants contributed at least one complete session, and 23 completed three or more complete sessions which was the threshold we required for within-person modelling. Among participants with at least one complete session, the median number of asm-EMA sessions was 7.5 (mean = 8.80, range = 1–27).

**Table 1.** Demographic information and clinical characteristics for enrolled subjects. Of 33 enrolled subjects, 30 went on to participate in the study. Duration of participation refers to the number of days that at least one asm-EMA session was triggered. Seizure type is calculated as the percentage of total patients who seized during hospitalization who experienced the listed type of seizure. These categories are not mutually exclusive; most patients experienced more than one type of seizure. *Of note, all 21 patients who seized experienced a spontaneous seizure, and 5 of the 21 patients who seized also experienced at least one stimulation-provoked seizure.

|  | N | Percentage | Median | IQR | Range |
|---|---|---|---|---|---|
| **Age** | 33 |  | 34 | 19 | 46 |
| **Sex** |  |  |  |  |  |
| Female | 22 | 66.7 |  |  |  |
| Male | 10 | 30.3 |  |  |  |
| Nonbinary | 1 | 3 |  |  |  |
| **Duration of participation (days)** | 30 |  | 4 | 2.8 | 9 |
| **Confirmed epileptogenic seizures** |  |  |  |  |  |
| Yes | 21 | 70 |  |  |  |
| No | 9 | 30 |  |  |  |
| **Seizure localization** |  |  |  |  |  |
| Temporal | 14 | 66.7 |  |  |  |
| Extratemporal | 4 | 19 |  |  |  |
| Mixed | 3 | 14.3 |  |  |  |
| **Seizure type (% of total subjects who seized)** \* |  |  |  |  |  |
| Spontaneous | 21 | 100 |  |  |  |
| Focal impaired awareness | 13 | 61.9 |  |  |  |
| Focal to bilateral tonic-clonic | 10 | 47.6 |  |  |  |
| Subclinical | 8 | 38.1 |  |  |  |
| Focal aware | 3 | 14.3 |  |  |  |
| Generalized (absence) | 2 | 9.5 |  |  |  |
| Stimulation-provoked | 5 | 23.8 |  |  |  |
| Focal impaired awareness | 3 | 14.3 |  |  |  |
| Focal aware | 2 | 9.5 |  |  |  |
| Subclinical | 1 | 4.8 |  |  |  |
| **Baseline neuropsychological assessment scores** |  |  |  |  |  |
| QOLIE-31 | 29 |  | 48 | 38 | 72 |
| BDI-II | 30 |  | 17 | 20 | 38 |
| BAI | 30 |  | 13 | 10.5 | 41 |
| STAI state | 22 |  | 47.5 | 28.5 | 55 |
| STAI trait | 22 |  | 42.5 | 21.5 | 42 |
| SBSOD | 16 |  | 4.15 | 1.75 | 5.2 |
| **Follow-up neuropsychological assessment scores** |  |  |  |  |  |
| QOLIE-31 | 12 |  | 54 | 30.8 | 63 |
| BDI-II | 12 |  | 12.5 | 13.5 | 34 |
| BAI | 12 |  | 13.5 | 7 | 28 |
| STAI state | 12 |  | 47 | 16.5 | 49 |
| STAI trait | 12 |  | 42 | 18.5 | 33 |
| SBSOD | 12 |  | 3.9 | 1.5 | 4.8 |
| **EEG type** |  |  |  |  |  |
| Scalp | 20 | 66.7 |  |  |  |
| Intracranial | 10 | 33.3 |  |  |  |

Session completion varied across participants and hospitalization days (Figure 2C). Although many participants contributed sessions across multiple days of monitoring, session density generally reflected hospitalization duration and individual engagement, with fewer observations available at later relative days as participants completed monitoring or were discharged. The number of sessions completed per day ranged from 1 to 7 (Figure 2D), with 1 to 4 sessions per day being the most common, consistent with the 90 to 150-minute inter-session triggering window.

**Figure 1.**
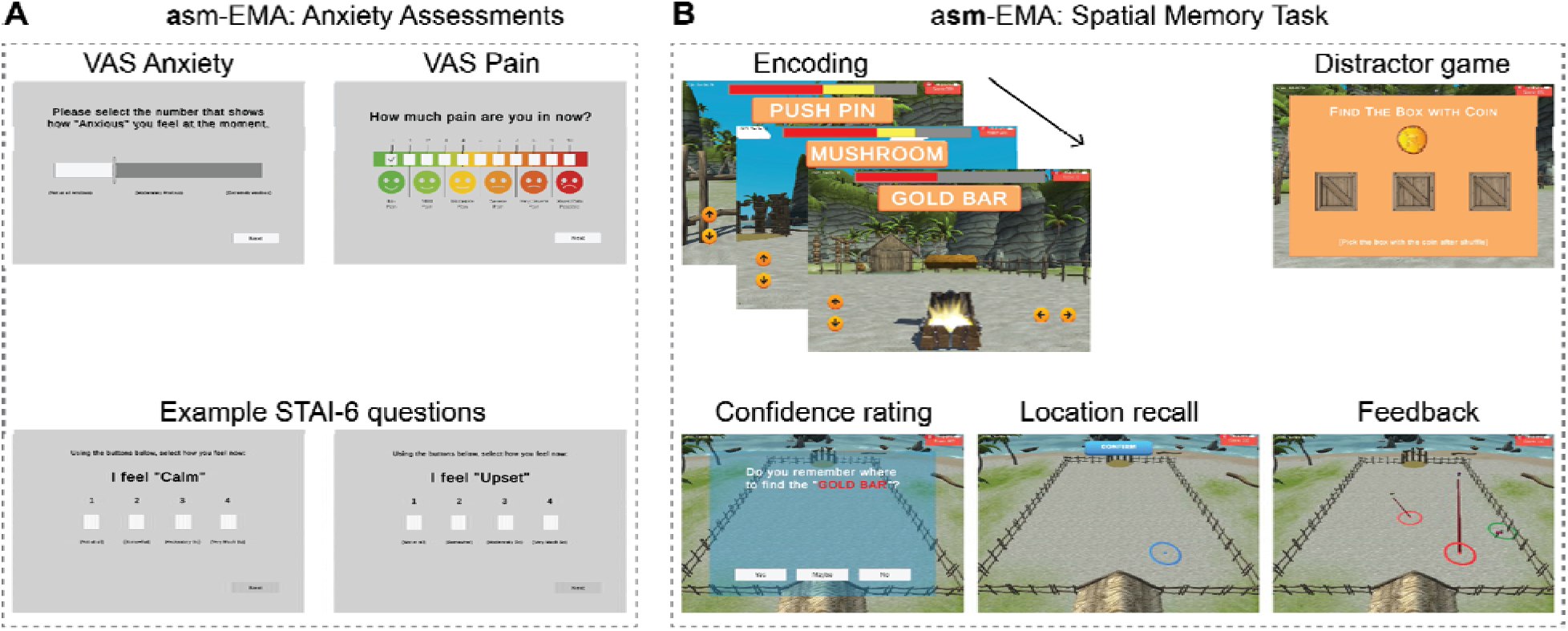
Structure of an anxiety-spatial-memory ecological momentary assessment (asm-EMA) session. Each asm-EMA session comprised two components delivered sequentially on a guided-access tablet. (A) Anxiety assessment. Participants completed a visual analog scale (VAS) for anxiety (top left), a VAS for pain (top right), and six items from the abbreviated State-Trait Anxiety Inventory (STAI-6; bottom), with example items shown for an anxiety-present (“I feel Upset”) and an anxiety-absent (“I feel Calm”) question. (B) Treasure Hunt spatial episodic memory task. In the encoding phase, participants navigated a virtual beach arena to locate treasure chests, each revealing an object (e.g., Push Pin, Mushroom, Gold Bar). An intervening distractor phase required tracking a coin shuffled between boxes. During the retrieval phase, participants indicated their confidence in remembering each object’s location before tapping the recalled position; corrective feedback displayed both the chosen (blue circle) and correct (red circle) locations.

**Figure 2.**
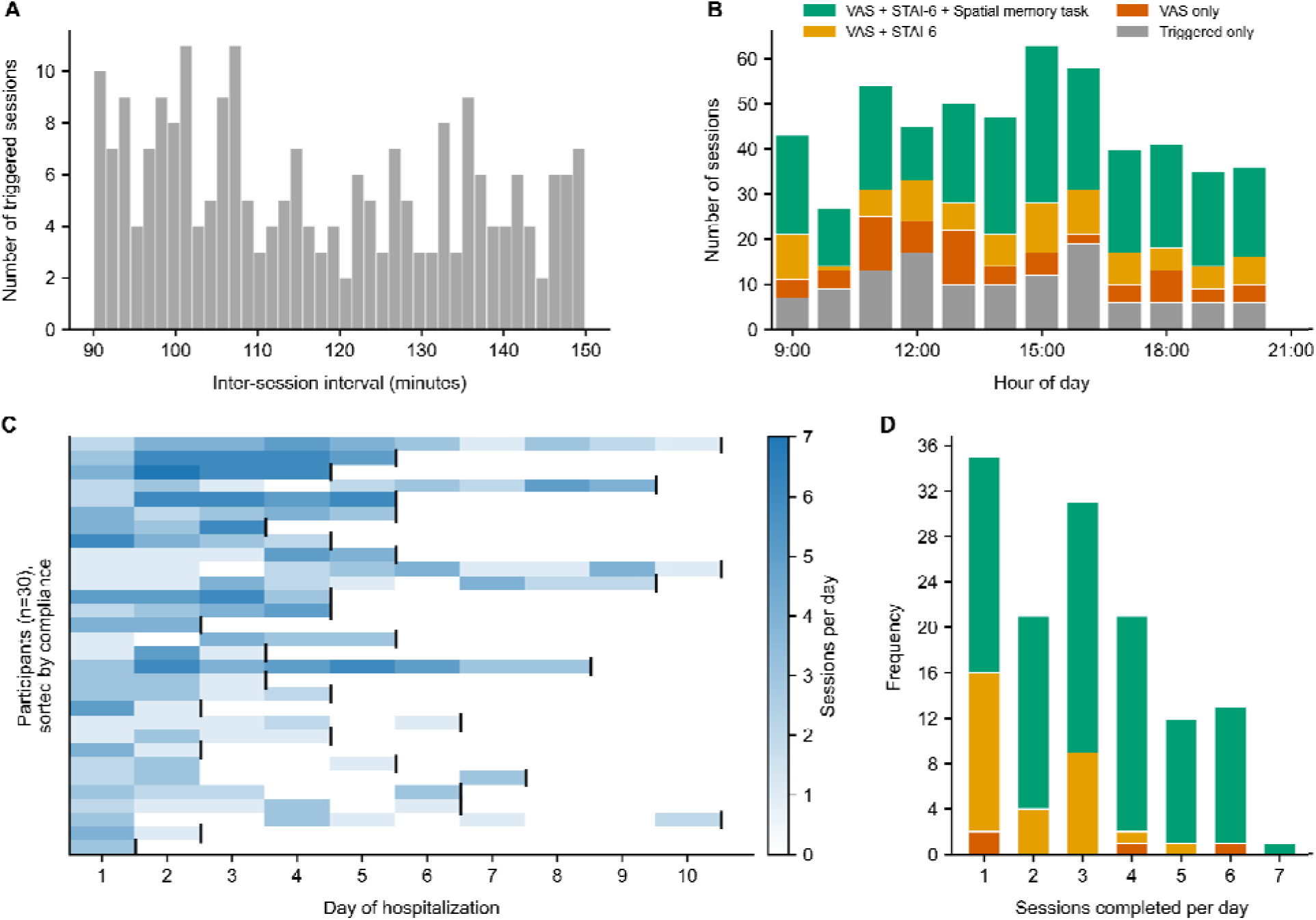
**asm**-**EMA protocol compliance and session delivery characteristics.** (A) Distribution of within-day inter-session intervals within the 90–150-minute triggering window. (B) Session completion by hour of day (09:00–21:00 EST), colored by completion tier: full protocol (VAS + STAI-6 + spatial memory task; green), full anxiety assessment (VAS + STAI-6; yellow), anxiety rating only (VAS only; orange), and triggered but not initiated (gray). (C) Session density per participant (rows, n = 30; sorted by total complete sessions) across relative hospitalization days (columns). White = no session; dark blue = highest daily session count. Vertical black lines indicate the end of the monitoring period. (D) Frequency of subject-days by number of sessions completed, colored by highest tier reached that day.

Session completion was distributed across 9am–9pm (Figure 2B). The full protocol constituted the largest category at nearly every hour of the day, indicating that time of day did not differentially drive dropout from the spatial memory component. Triggered-only sessions, in which prompts that were delivered but not initiated, were present throughout the day but did not cluster at any particular timepoint.

### asm-EMA Protocol Validity and Variability

Subject- and session-level distributions for all five asm-EMA measures are shown in Figure 3. The two momentary anxiety measures showed markedly different distributions. VAS Anxiety was strongly right skewed at the session level, with ratings concentrated near zero and a long tail extending to the scale maximum (Figure 3B), consistent with an inpatient sample in which acute anxiety is episodic rather than sustained. STAI-6 scores were more broadly distributed across the 6 to 24 range, with subject-level means spanning much of the scale (Figure 3C). This difference likely reflects measurement format where the single-item VAS is sensitive to acute distress but compresses responding at the low end, whereas the multi-item STAI-6 distributes variance more evenly by aggregating across items with differing affective content. VAS Pain showed the most pronounced floor effect of the three self-reported measures, with the majority of sessions rated at or near zero while a subset maintained persistently elevated pain levels throughout hospitalization (Figure 3A). Memory accuracy was approximately normally distributed at the session level, centered near 0.75, indicating performance consistently well above chance (Figure 3D). Log-transformed response times were approximately unimodal, with subject-level means clustering around the center of the session-level distribution (Figure 3E). Variance decomposition from random-intercept models indicated that all five asm-EMA measures contained both stable between-subject differences and within-subject session-to-session variation. Between-subject differences accounted for 53%, 47%, 66%, 54%, and 69% of total variance in VAS Pain, VAS Anxiety, STAI-6, memory accuracy, and log-transformed response time, respectively (Figure 3F). Bootstrap confidence intervals around the ICC estimates indicated that these variance-decomposition estimates were reasonably stable, while also reflecting uncertainty due to the modest sample size and unequal number of sessions per participant. VAS Anxiety showed the largest within-subject proportion, with 53% of variance occurring within individuals, despite its floor-skewed distribution. STAI-6 showed lower within-subject variance, with 34% of variance occurring within individuals, consistent with item-level averaging dampening momentary reactivity relative to a single-item format. Across all five measures, within-subject variance ranged from 31% to 53%, supporting the use of repeated asm-EMA measurements to examine session-level anxiety-memory covariation.

**Figure 3.**
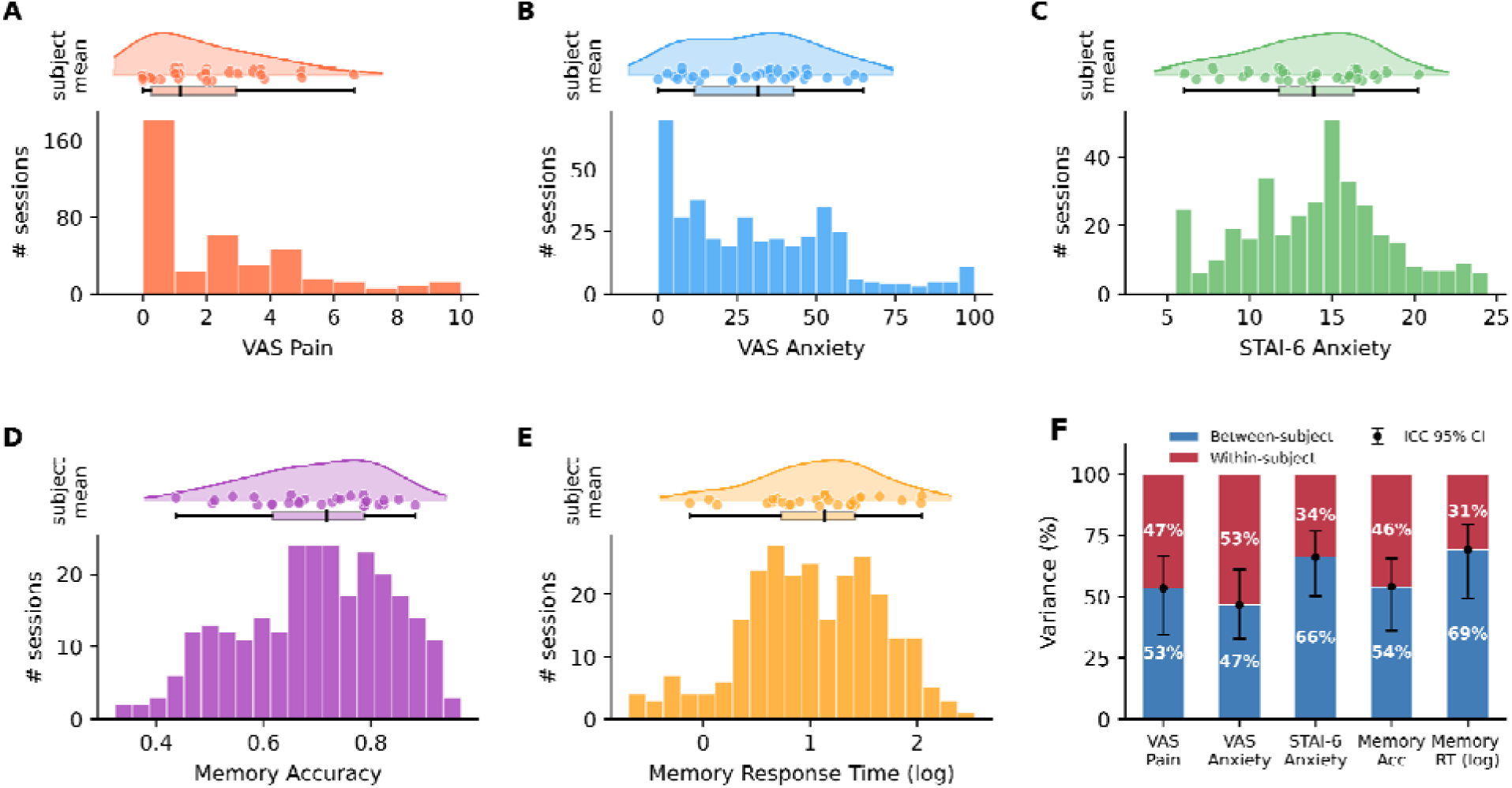
Session-level distributions and subject-level variability for all asm-EMA measures. (A-E): each shows two views of the same measure. The upper portion displays a raincloud plot of subject-level means, comprising a kernel density estimate, individual subject data points (jittered for visibility), and a box plot (median, IQR, and 1.5× IQR whiskers). The lower portion shows the session-level frequency distribution across all valid asm-EMA sessions. Measures shown are VAS Pain (A), VAS Anxiety (B), STAI-6 (C), Treasure Hunt memory accuracy (D), and Treasure Hunt logged response time (E). (F): variance decomposition for each asm-EMA measure, estimated from random-intercept linear mixed-effects models. Stacked bars show the proportion of total variance attributable to between-subject differences and within-subject session-to-session variation. Points and error bars indicate ICC estimates with 95% bootstrap confidence intervals.

### asm-EMA Session Mean and Variability Align with Standard Neuropsychological Assessments

Subject-level asm-EMA session means tracked standard neuropsychological assessments collected before and after the monitoring period (Figure 4A; Figure S2A; scatter plots of all significant correlation pairs in Figure S4). STAI-6 showed the strongest and broadest correlation with the full STAI State and Trait inventories at ρ = 0.80–0.85 (Figure 4C, right panel), with BAI (ρ = 0.54, p < .01) and BDI-II (ρ = 0.61, p < .001), and negatively with QOLIE-31 (ρ = −0.62, p < .001). The average VAS Anxiety and VAS Pain showed broadly similar positive correlations with BAI (ρ = 0.51, p < .006 for both) and STAI State (VAS Anxiety: ρ = 0.49, p < .027; VAS Pain: ρ = 0.46, p < .044). Among the memory measures, response time correlated with BAI (ρ = 0.40, p < .05), suggesting more anxious individuals taking longer to commit during retrieval. The trending positive correlation between spatial memory accuracy and the Santa Barbara Sense of Direction scale (SBSOD, ρ = 0.48, p < .078) suggests the task reflects genuine spatial cognitive ability.

**Figure 4.**
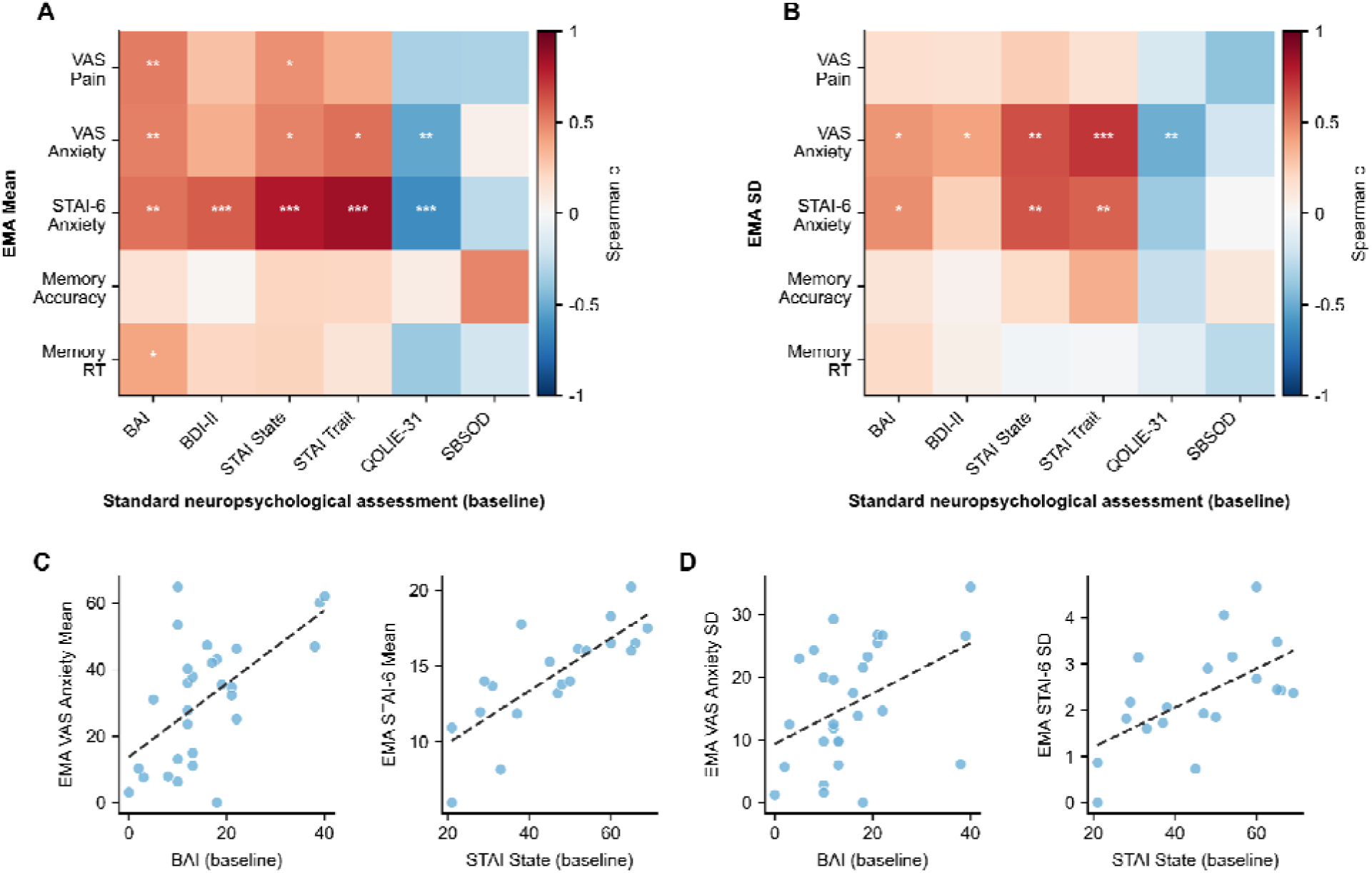
Convergent validity of asm-EMA means and within-person variability against standard neuropsychological assessments. Heatmaps show Spearman correlations between subject-level asm-EMA summary statistics (rows) and standard clinical measures administered at admission (columns): BAI, Beck Anxiety Inventory; BDI-II, Beck Depression Inventory; STAI State and Trait, State-Trait Anxiety Inventory; QOLIE, Quality of Life in Epilepsy; SBSOD, Santa Barbara Sense of Direction scale. Color encodes correlation magnitude and direction (red = positive, blue = negative); asterisks denote significance (* p < .05, ** p < .01, *** p < .001). (A) Correlations with asm-EMA subject means. (B) Correlations with asm-EMA within-person standard deviations. (C, D) Scatter plots illustrate representative associations between admission BAI and STAI State scores (x-axes), and asm-EMA VAS Anxiety mean and STAI-6 mean (C) and their within-person standard deviations (D). Dashed lines show ordinary least-squares fits.

Within-person variability (measured as standard deviation; SD) was also systematically predicted by baseline clinical severity (Figure 4B). VAS Anxiety SD showed the broadest pattern of associations, correlating significantly with BAI (Figure 4D, left panel), BDI-II, STAI State, STAI Trait, and QOLIE, indicating that patients with more severe baseline psychopathology fluctuated more from session to session. STAI-6 SD was more selective, associating significantly with STAI State (Figure 4D, right panel) and STAI Trait but not with BDI-II or QOLIE, consistent with it indexing state anxiety reactivity more specifically than general affective severity. Neither memory measure nor VAS Pain showed significant SD associations with clinical instruments, suggesting that moment-to-moment fluctuations in pain and memory are driven more by proximal situational factors than by trait-level severity. The correspondence between asm-EMA measures and standard clinical instruments was maintained at follow-up, though the reduced sample at that timepoint limits the interpretive weight of those replications (Figure S2; scatter plots of all significant correlation pairs in Figure S5). Additional sensitivity analyses using a minimum threshold of five valid observations and observation-weighted rank correlations showed a similar pattern, and FDR-adjusted q-values are reported to account for multiple comparisons across the clinical validity correlations (Figure S3).

### Covariation Between Momentary Anxiety State and Spatial Memory Performance

We computed the between-subject correlations using subject-level means, and within-subject correlations using person-mean-centered session-level observations. At both levels, VAS Pain, VAS Anxiety, and STAI-6 were positively correlated with one another, supporting convergence among the affective measures (Figure 5A, B; Figure S6). Memory accuracy showed no reliable association with momentary anxiety or pain at either level. In contrast, within-subject correlations showed a negative association between STAI-6 and memory response time, indicating that sessions with higher-than-usual STAI-6 anxiety were associated with faster responses (ρ = -0.154, p < 0.013; Figure 5B).

**Figure 5.**
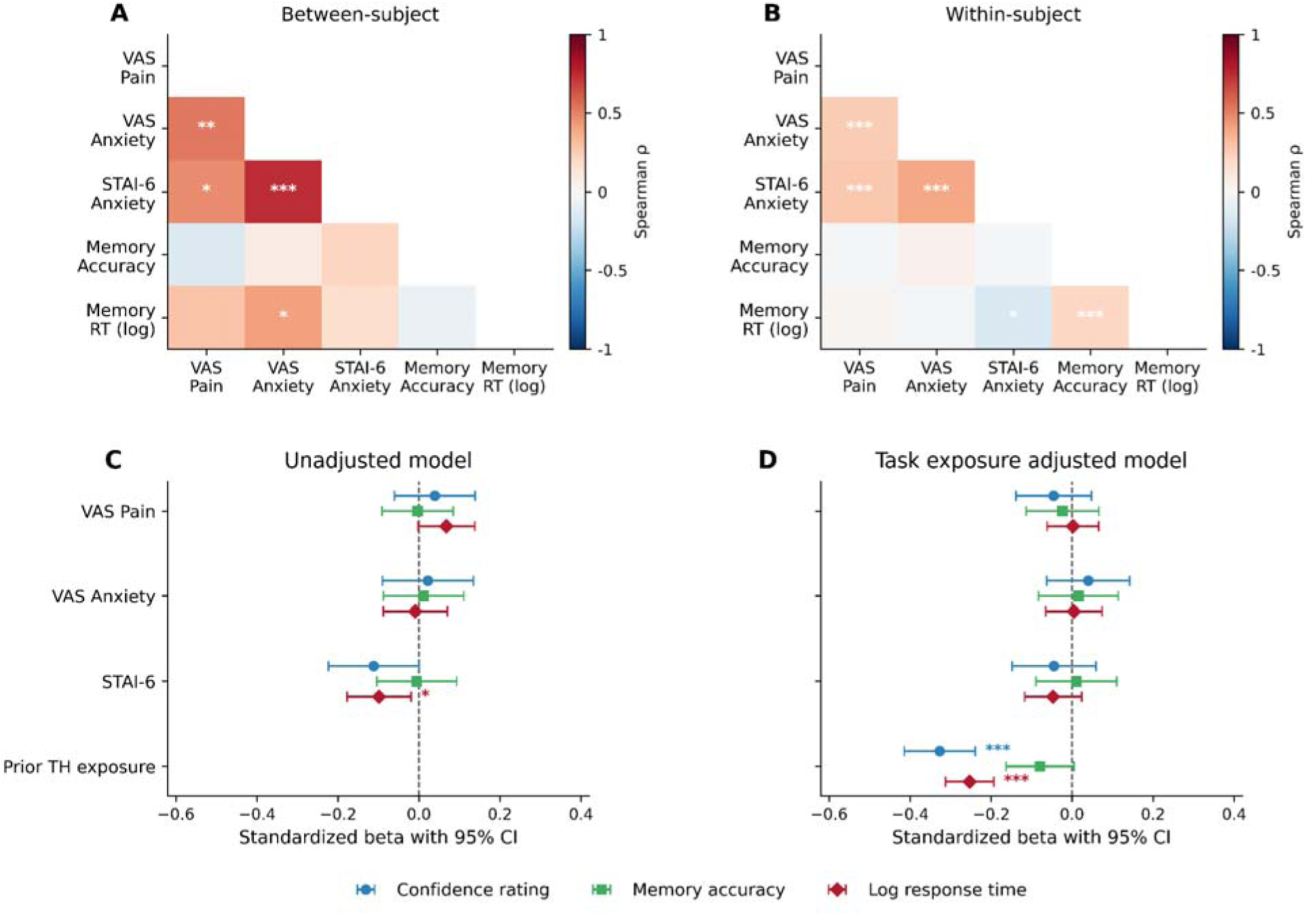
Within- and between-person associations among asm-EMA measures and their relationship to memory performance. Panels A and B show Spearman correlation matrices for between-subject and within-subject associations, respectively. (A) displays correlations computed on subject-level means; (B) displays correlations computed on person-mean centered session-level data. In both panels, only the lower triangle is shown and asterisks indicate significance (* p < .05, ** p < .01, *** p < .001); color encodes the Spearman ρ value. (C) Unadjusted multivariable mixed-effects models testing concurrent associations between VAS Pain, VAS Anxiety, STAI-6, and task performance. (D) Exposure-controlled models additionally including prior Treasure Hunt exposure, defined as the number of previously completed Treasure Hunt sessions. Points indicate standardized beta coefficients and error bars indicate 95% confidence intervals. Significance markers indicate p < 0.05, p < 0.01, and p < 0.001.

We next used multivariable mixed-effects models to examine concurrent associations between momentary anxiety state and the two primary spatial memory outcomes, while accounting for repeated observations within participants. Normalized memory accuracy and log-transformed response time were modeled separately. In the original models, VAS Pain, VAS Anxiety, and STAI-6, were not associated with memory accuracy. In contrast, higher STAI-6 was associated with faster log response time (beta = -0.098, 95% CI [-0.177, -0.020], p < 0.015), whereas VAS Anxiety and VAS Pain were not significant predictors (Figure 5C).

To account for potential familiarity with the spatial memory task across repeated sessions, we then repeated the models with prior task exposure included as a covariate. Memory accuracy remained unrelated to the VAS Pain, VAS Anxiety, and STAI-6 ratings. For response time, prior task exposure was strongly associated with faster responses (beta = -0.253, 95% CI [-0.313, -0.194], p < 0.001). After accounting for prior task exposure, the association between STAI-6 and log response time was attenuated and no longer statistically significant (beta = -0.047, 95% CI [-0.117, 0.023], p = 0.192; Figure 5D). A useful direction for future work is to understand the relation between memory and anxiety while accounting for practice and time effects.

As a complementary task-derived measure, we also examined confidence ratings collected during the Treasure Hunt retrieval phase. Confidence ratings showed expected associations with task behavior, such that higher confidence was associated with lower spatial error, higher normalized accuracy, and longer response times (Figure S7). In the original mixed-effects model, affective predictors were not significantly associated with confidence, although higher STAI-6 was associated with lower confidence (beta = -0.111, 95% CI [-0.223, 0.001], p = 0.051; Figure 5C-D). Additional sensitivity analyses showed that including EEG type as a covariate did not materially alter the primary mixed-effects model estimates, indicating that the anxiety-memory findings were not driven by differences between SEEG and scalp EEG participants (Figure S8, Figure S9).

### Short-Term Temporal Persistence and Predictability of Anxiety and Memory Performance from asm-EMA

We next examined whether deviations from each participant’s average asm-EMA level were carried forward from one session to the next. Under our sampling protocol of approximately 2-hour intervals, this allowed us to characterize short-term dynamics between consecutive assessments and evaluate whether the sampling frequency captured meaningful within-person change. We assessed these dynamics using within-person Spearman autocorrelations (see methods), which tested whether person-relative values of the same measure were correlated across adjacent and later sessions (excluding cross-day transitions and unusually long gaps). Significance was evaluated using permutation testing (Figure 6A), and we found significant positive autocorrelation between adjacent sessions (lag one) in four of five measures: VAS Pain (ρ = 0.31, p < .01), VAS Anxiety (ρ = 0.29, p < .01), STAI-6 Anxiety (ρ = 0.25, p < .01), and memory RT (ρ = 0.37, p < .001), with spatial memory accuracy showing a near-significant lag-one effect (ρ = 0.18, p < .10). Beyond lag one, autocorrelations were largely attenuated for the anxiety measures and spatial memory accuracy (Figure 6A; Figure S10), indicating that temporal dependence was concentrated at the immediately preceding session. Response time was the exception, with significant persistence extending through lag three. Together, these lag profiles indicate statistically significant but modest short-term persistence in person-relative state deviations, with little evidence that this dependence extended to longer lags. Because these correlations were modest rather than large, repeated assessments appeared to capture related but not unchanged states.

**Figure 6.**
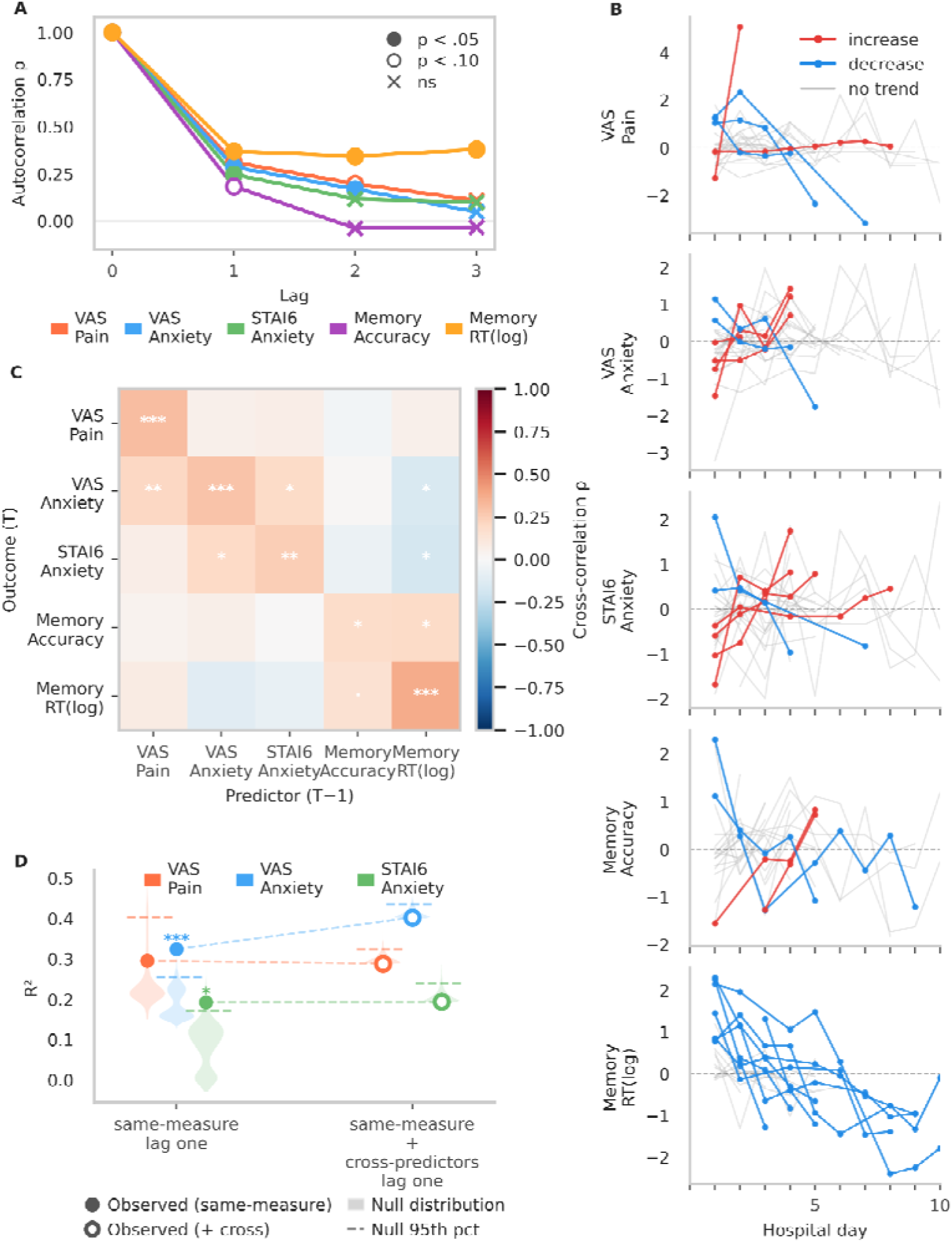
Temporal structure of asm-EMA measures. (A) Pooled within-subject Spearman autocorrelation (ρ) at lags 0–3 for all five measures. Filled circles indicate p < .05, open circles p < .10, and crosses non-significant, based on permutation testing. (B) Within-patient trajectories across hospital days for each measure. Each line represents one patient; points show daily means. Lines are colored by the direction and significance of a within-patient linear trend fitted on hospital day: red = significant increase, blue = significant decrease, grey = non-significant. (C) Lag-1 cross-correlation matrix (pooled Spearman ρ); rows = outcome at T, columns = predictor at T−1. Asterisks denote permutation significance (· p < .10, * p < .05, ** p < .01, ***< .001). (D) Permutation-tested predictability of the affective autoregressive and cross-predictor models. Filled circles show the observed R² for same-measure lag-one autoregressive models (left), testing whether the previous-session value improved prediction beyond the covariates. Open circles show the observed R² after adding lag-one values of the other affective measures as cross-predictors (right), testing whether they improved prediction beyond the same-measure lag-one term and covariates. Shaded areas show the corresponding within-subject permutation null distributions, dashed lines mark their 95th percentiles, and asterisks indicate permutation significance.

To further characterize temporal structure across different timescales, we first examined within-subject trajectories across hospital days. A subset of participants showed significant increases or decreases over hospitalization (Figure 6B), indicating that gradual hospital-day trends represented a structured component of temporal variation rather than merely a nuisance source of adjacent-session correlation. We quantified this structure by modeling each measure as a function of hospital day with participant-specific intercepts and slopes and evaluating the explained variance against a within-subject permutation null (Table 2). Hospital day explained the most variation in Memory RT (R² = .365, p < .001), with trajectories predominantly decreasing, consistent with increasing task familiarity. Among the anxiety measures, hospital day explained more variation in STAI-6 Anxiety (R² = .147, p < .001) than in VAS Anxiety (R² = .050, p = .014), indicating greater gradual hospitalization-related structure in STAI-6.

**Table 2.** Daily trend and autoregressive self-prediction at lag-1. Day-Trend R² is the proportion of total within-person variance explained by a linear mixed-effects model with hospital day as the fixed predictor and participant-specific intercepts and slopes; p is the one-tailed permutation p-value (proportion of 2,000 null R² values ≥ observed R²). AR R² summarizes model performance for the lag-1 autoregressive model fitted to detrended scores (detrended lag-1 score, elapsed time, time of day, session number; random intercept and random lag-1 slope per participant), evaluated against a within-participant permutation null generated by shuffling the detrended lag-1 predictor. β lag-1 is the fixed-effect estimate for the lag-1 predictor from the mixed-effects model, with 95% confidence interval and analytic p-value. Bold highlights statistical significance.

| asm-EMA measures | Day Trend **R<sup>2</sup>** | Day trend p (perm) | Detrend-AR R<sup>2</sup> | Detrend-AR p (perm) | Detrend-AR lag-1 β | 95% CI | Detrend-AR lag-1 p |
|---|---|---|---|---|---|---|---|
| VAS Pain | 0.041 | 0.051 | 0.295 | 0.161 | +0.016 | [-0.135, +0.168] | 0.831 |
| VAS Anxiety | 0.050 | **0.014** | 0.324 | **0.000** | +0.182 | [+0.007, +0.357] | **0.042** |
| STAI-6 Anxiety | 0.147 | **0.000** | 0.192 | **0.026** | +0.062 | [-0.186, +0.309] | 0.626 |
| Memory Accuracy | 0.153 | **0.001** | 0.185 | 0.645 | -0.019 | [-0.124, +0.087] | 0.726 |
| Memory RT (log) | 0.365 | **0.000** | 0.219 | **0.059** | -0.088 | [-0.317, +0.141] | 0.449 |

We next removed each subject’s linear hospital-day trajectory and fitted mixed-effects autoregressive models to the detrended values. These models tested whether knowing the previous-session value improved prediction beyond elapsed time, time of day (with modest differences between morning and afternoon sessions; Figure S11), session number, and implant type. We used permutation-tested R² to assess added predictive value and the lag-one coefficient to characterize the direction of the association (Fig. 6D, left; Table 2, Table S1, Table S2). Prior-session predictability was stronger and directionally consistent for VAS Anxiety (R² = .324, p < .001; β = .182, 95% CI [.007, .357], p = .042) than for STAI-6 (R² = .192, p = .026; β = .062, 95% CI [−.186, .309], p = .626). The stronger residual predictability of VAS Anxiety after detrending suggests that removing hospital-day trends had less impact on its short-term persistence, consistent with its weaker hospitalization-related structure relative to STAI-6. Beyond the lagged effects, the same models showed that later time of day was associated with lower Memory Accuracy (β = −.134, p = .018; Figure S12), potentially reflecting fatigue or other accumulating within-day influences on memory performance.

To test whether person-relative fluctuations in one domain were followed by changes in another, we examined cross-correlations among all five asm-EMA measures, asking whether person-relative deviations in one measure at a given session were correlated with person-relative deviations in another measure at a later session (permutation testing; see methods). Some anxiety measures showed significant cross-correlations with different measures at the following session (lag one); for example, higher-than-usual VAS Pain was correlated with higher-than-usual VAS Anxiety at the following session (ρ = .21, p < .01; Figure 6C). Cross-correlations at longer lags are shown in Figure S13. These findings suggest possible short-term coupling between measures. However, cross-correlation alone cannot tell whether one measure contributes predictive information about later person-relative deviations in another, because an apparent lagged correlation may arise when each measure is persistent over time and the two measures are also correlated within the same session. We therefore extended the autoregressive models by adding lagged predictors from the other affective measures, referred to here as cross-predictors. For each anxiety outcome, we tested whether these cross-predictors improved prediction beyond the same-measure lag-one predictor. No significant cross-predictor effects were found for any measures (Figure 6D). Adding the other affective measures from the previous session did not significantly improve model performance beyond the same-measure lag-one predictor (all p > .82 for all outcomes; within-subject permutation testing, details in methods). These results indicate that cross-correlations between different measures did not add predictive information once each outcome’s own previous-session person-relative value was accounted for. Thus, the main temporal pattern in the asm-EMA data was persistence of values within the same measure across adjacent sessions.

## Discussion

A central motivation of this study was to close the ecological validity gap in anxiety–memory research by measuring mood and memory together, repeatedly, and in context. The asm-EMA protocol is well positioned to this goal because it captures subject state in near-real-time in natural environments, reducing dependence on retrospective report and improving ecological validity^15,31^. Prior naturalistic cognitive studies have established that brief smartphone-based tasks can be used outside the laboratory to obtain reliable cognitive measurements and to track meaningful within-person cognitive variation over time^32,22^. The present findings extend that evidence to a clinical inpatient setting. The asm-EMA protocol produced interpretable, moment-to-moment measures of anxiety and spatial memory across multiple days of hospitalization, identifying substantial within-person variability in both domains.

Subject-level asm-EMA means showed strong correspondence with the baseline and follow-up standard clinical assessments. The strong association between asm-EMA STAI-6 scores and the full STAI supports the convergent validity of the abbreviated measure in this inpatient EMA context. These findings suggest that STAI-6 can efficiently capture clinically meaningful variation in anxiety during repeated monitoring, while not establishing formal equivalence with the full inventory. Spatial memory accuracy correlated positively with the SBSOD scale, confirming that the task indexes a distinct spatial cognitive ability rather than a proxy for affective state. Beyond mean level, asm-EMA protocol captured a second clinically meaningful dimension which is the within-person variability. Patients with higher baseline clinical anxiety showed greater moment-to-moment STAI-6 fluctuation during hospitalization, indicating that trait severity predicts not just where someone resides on the anxiety scale on average, but how much they fluctuate over time. A single clinical snapshot misses this dimension entirely. The correspondence between asm-EMA means and standard neuropsychological instruments was largely maintained at follow-up, though the reduced sample at that timepoint limits the interpretive weight of these replications.

Importantly, there was no evidence of response bias on the basis of anxiety level at the time of the EMA alert. If patients with greater anxiety were less likely to complete the full asm-EMA session, the anxiety scores in subjects completing the more extensive memory assessment would be systematically lower, biasing any anxiety-memory comparison. The data do not support this systematic dropout bias. Brief VAS Anxiety and VAS Pain assessments, mandatory to silence the alert, were completed after 85.5% of alerts, did not differ based on the extent of asm-EMA session completion (VAS Anxiety: Kruskal-Wallis H = 1.49, p > .474; VAS Pain: H = 2.02, p > .364; Figure S1), with group means within 3 points on a 100-point scale. Within-session dropout was effectively independent of anxiety state at onset.

Across analytic levels, we did not detect evidence that anxiety was associated with reduced spatial memory accuracy. Participants with higher average anxiety did not show lower average memory accuracy, and sessions with higher-than-usual anxiety were not associated with lower memory accuracy within the same participant. These findings extend prior laboratory work reporting anxiety-related memory impairment under controlled stress conditions^5,6^ to a naturalistic clinical context, where the relationship may operate differently. Whether this reflects genuine memory resilience under ecological conditions, adaptation to chronic stress, or anxiety levels below the threshold for memory disruption remains an open question worth pursuing in future work.

The response-time findings revealed a potentially informative dissociation between memory accuracy and retrieval behavior. In the original within-person mixed-effects model, higher STAI-6 was associated with faster log-transformed retrieval responses, even after adjusting for VAS Anxiety and VAS Pain, while memory accuracy remained unchanged. This pattern is consistent with the distinction between performance effectiveness and processing efficiency proposed in attentional control and processing efficiency frameworks^1,33^, in which anxiety may alter the time or effort devoted to retrieval without necessarily changing accuracy. Importantly, however, the response-time association was attenuated after controlling for prior task exposure, which was itself strongly associated with faster responding. Supplemental analyses further showed that momentary anxiety was not robustly associated with confidence ratings or continuous spatial error. Thus, the response-time effect should not be interpreted as direct evidence for a specific retrieval strategy, but it does suggest that response timing may be a sensitive behavioral dimension for detecting state-dependent changes during repeated cognitive assessment.

Our temporal analyses suggest that the asm-EMA protocol captured structured within-subject variation at both short and longer timescales. In the emotion-dynamics literature, lag-one autoregressive carryover is often interpreted as emotional inertia, or the extent to which an affective state persists into the next measurement rather than changing immediately^34^. More recent EMA-based work has extended this framework by allowing the strength and direction of affective interactions to vary across days within an individual^35^. Consistent with this broader framework, our asm-EMA protocol captured a pattern of short-lived inertia, with associations strongest between adjacent sessions and largely decaying at later lags. Beyond this short-term persistence, several measures also showed structured change across the duration of the hospitalization. Such patient-specific trajectories may contain clinically meaningful information that is not visible in aggregate scores and may support a more individualized understanding of symptom course^36^. In our study, this slower structure was stronger for STAI-6 than for VAS Anxiety, suggesting that these components of asm-EMA capture both momentary continuity and broader changes in patients’ affective states over an inpatient stay.

A practical question raised by these findings is whether the roughly two-hour sampling interval was appropriate for capturing within-person state variation. The same schedule revealed different temporal profiles even for two measures of anxiety: VAS Anxiety showed clearer session-to-session persistence, whereas STAI-6 changed more gradually across hospitalization. Prior work similarly notes that different state-anxiety instruments may have separate theoretical bases, item content, format, and sensitivity and may yield different findings despite targeting the same broad construct^37^. Thus, even assessments targeting the same construct or cognitive function may require different sampling schedules. More broadly, less frequent sampling may be sufficient for questions about broad daily state, whereas denser sampling may be needed to distinguish fine-grained changes following a seizure, medication change, procedure, or other clinical event. A strength of the asm-EMA framework is that its automated scheduling system can be adapted to these different temporal scales, allowing future studies to tailor the assessment battery and sampling interval to particular scientific or clinical questions.

To our knowledge, this is the first study to combine repeated ecological momentary anxiety assessment with a performance-based spatial memory task in a clinical inpatient population, and the first to characterize within-person anxiety-memory covariation under naturalistic clinical stress conditions. Prior EMA work in cognitive domains has relied almost exclusively on community samples and brief attentional tasks^22,23,25,26^. Extending this methodology to an inpatient epilepsy setting with a validated episodic memory paradigm represents a substantive step toward ecologically grounded cognitive monitoring in clinical practice.

These findings have several practical implications for inpatient EMU assessment. Across analyses, spatial memory accuracy was not systematically reduced during periods of elevated anxiety or pain, suggesting that it was relatively stable across the affective fluctuations observed in this sample. This pattern is reassuring for repeated cognitive assessment during EMU admission, although it should not be interpreted as evidence that anxiety is unrelated to memory under all clinical conditions. Response time appeared more strongly related to prior task exposure than to momentary anxiety state after accounting for repeated assessments, indicating that faster responding across hospitalization may reflect increasing task familiarity rather than improved memory performance or a specific anxiety-related retrieval strategy. Finally, associations between asm-EMA anxiety summaries and standard clinical inventories support the convergent validity of brief repeated anxiety assessments in this setting. These results suggest that intensive short-form assessments may help monitor anxiety burden and affect variability during hospitalization while complementing standard clinical measures.

Several limitations warrant consideration. The cohort was modest in size and protocol completion was uneven across participants, limiting statistical power for weaker effects and confidence in subgroup-level inference. The anxiety-memory findings are also conditional on task completion. Because participants could skip EMA prompts or stop after the self-report portion, the cognitive analyses may under-sample periods of high distress, fatigue, disengagement, or clinical instability. VAS anxiety and pain scores did not differ significantly across completion tiers, providing some reassurance against systematic bias, though subtler forms of selective missingness cannot be ruled out completely. Consistent with this, a session-level sensitivity analysis showed that VAS Anxiety and STAI-6 did not predict Treasure Hunt completion, although higher VAS Pain and later session number were associated with modestly lower odds of completion (Figure S1). The present analysis also focused on average within-person temporal patterns rather than individual differences in affective dynamics. Prior work suggests that affective inertia varies substantially across individuals and is elevated in those with depressive symptoms^34^, and characterizing this heterogeneity in future studies would enrich the picture considerably.

The clinical context in which our study was conducted introduces additional confounds that cannot be fully resolved here. The prompting procedure in the inpatient EMU setting may have transiently increased variability in VAS Anxiety, which was assessed while the alert remained active, whereas STAI-6 was completed after the alert had stopped and was therefore less directly affected by the prompt. Patients with epilepsy show elevated rates of anxiety and memory impairment relative to the general population^38^, and those with temporal lobe involvement may show altered anxiety-memory dynamics specifically^39^. Antiseizure medication withdrawal, standard practice during EMU monitoring, may independently modulate both anxiety and cognition across the monitoring period. Seizure timing relative to individual sessions was not systematically tracked, and post-ictal confounding of affective and cognitive ratings in a subset of observations cannot be excluded. Prospective studies should define and separately analyze sessions falling within a post-ictal window, as these states may differ substantially from interictal baseline.

Our study represents a foundational step within the broader Context-Aware Multimodal Ecological Research and Assessment platform^30^, which aims to build a context-aware framework for tracking internal state using synchronized multimodal data including intracranial recordings, physiological monitoring, and passively acquired behavioral signals. Within that framework, the asm-EMA protocol serves two functions, as a standalone clinical assessment tool and as a source of structured behavioral ground truth for predictive modeling. The short-term temporal dependence observed between adjacent sessions is particularly relevant for the latter, since asm-EMA labels preserve local continuity rather than acting as isolated snapshots and provide a natural structure for training state-prediction models that learn from shared short-term context without redundancy. As the platform scales to larger samples and richer data modalities, the behavioral and clinical validity established here provides the empirical foundation on which multimodal predictive models can be built and interpreted.

## Methods

Our study used anxiety-spatial-memory ecological momentary assessment (asm-EMA) protocol to test the relationship between momentary anxiety and spatial episodic memory in epilepsy patients undergoing EEG monitoring. Participants completed repeated asm-EMA sessions throughout their hospitalization, each comprising a brief anxiety assessment battery followed by a spatial memory task. Sessions were delivered automatically at pseudo-randomized intervals across the day, allowing us to capture naturalistic fluctuations in anxiety state and cognitive performance within and across individuals.

### Recruitment and Enrollment

Potential subjects were identified from patients with known or suspected epilepsy scheduled for admission to the New York-Presbyterian/Columbia University Irving Medical Center EMU for non-invasive scalp video electroencephalography (EEG) or intracranial video stereoelectroencephalography (SEEG). Preliminary screening was conducted via electronic medical record review. Eligible subjects were aged 18–65, of any gender, proficient in English or Spanish, and had at least a 7th-grade reading level. Exclusion criteria included significant hearing, visual, or intellectual impairment; history of electroconvulsive therapy or psychosis (excluding postictal psychosis); lifetime history of psychotic disorder; current severe or unstable anxiety, major depressive disorder, or bipolar disorder; neurodegenerative disease or widespread brain lesions; language impairment; or medical conditions potentially affecting cognition (HIV, metastatic cancer, acute renal failure, or end-stage renal disease).

Once a subject was deemed eligible, the research team notified the treating physician to confirm appropriateness for enrollment. SEEG patients were contacted by phone prior to admission to introduce the study, while scalp EEG patients were approached on the day of admission in the EMU. Formal written informed consent was obtained from all participants in person, typically on the day of admission. The study was conducted in accordance with the Declaration of Helsinki and approved by the Columbia University Irving Medical Center Institutional Review Board, Columbia University, New York, NY, USA, approval number AAAU7374.

### Anxiety-Spatial Memory Ecological Momentary Assessment (asm-EMA)

During each asm-EMA session, patients were first prompted to report what they were doing immediately prior to the alert from a pre-specific multiple-choice list of activities. They were next prompted to complete three self-report instruments assessing current anxiety and pain. State anxiety was measured using a visual analog scale (VAS Anxiety), presented as a horizontal slider anchored at “Not at all anxious” (0) and “Extremely Anxious” (100). Current pain was measured with a similar VAS (VAS Pain), anchored at “No Pain” (0) and “Worst Pain Possible” (10), with emoticons accompanying the scale to aid interpretation. Both scales required participants to move a slider to indicate their current state (Figure 1A top row).

Patients then completed the STAI-6 (Figure 1A bottom row), a validated six-item abbreviation of the Spielberger State-Trait Anxiety Inventory designed for settings where the full 40-item form is impractical^40^. The six items were selected from the 20-item state anxiety subscale based on their correlation with the remaining items, comprising the three highest-loading anxiety-present and three highest-loading anxiety-absent items. Each item is rated 1–4, with anxiety-absent items reverse-scored before summation, yielding a total score ranging from 6 to 24 where higher scores indicate greater state anxiety. The STAI-6 has been validated against the full STAI through concurrent administration across populations with both average and elevated anxiety levels.

Following the anxiety assessment, participants completed the Treasure Hunt task, a spatial episodic memory paradigm involving virtual navigation, adapted from previous studies^28,29,41^. Using a touchscreen iPad, participants navigated a virtual beach arena (100 × 70 virtual units), encountered then later retrieved a series of treasure chests at randomized locations (Figure 1B).

Each trial consisted of three phases. In the encoding phase, four chests appeared one at a time, and participants navigated to each in sequence. Upon reaching a chest, it opened to reveal either an object (displayed for 1.5s) or an empty interior; two or three of the four chests contained objects, and participants were instructed to remember their locations. A brief distractor phase followed, in which participants tracked a coin hidden under one of three boxes as the boxes repeatedly swapped positions. During the retrieval phase, participants were presented with each encountered object in random order and asked to rate their confidence in remembering its location (“Yes,” “Maybe,” or “No”) before tapping the remembered location directly on the arena. Responses within 11 virtual units of the true location were displayed as correct, and responses outside this threshold were displayed as incorrect.

For statistical analyses, memory accuracy was analyzed as a continuous normalized measure rather than as the binary feedback outcome shown to participants. For each response, we calculated the Euclidean distance between the true object location and the selected response location. Raw distance errors were then converted to normalized accuracy scores based on their percentile rank relative to all possible response errors for that item location, following^29^. Scores ranged from 0 to 1, with 1 indicating a perfect response and 0 indicating the worst possible response. Session-level memory accuracy was computed as the mean normalized accuracy score across all attempted retrievals in that session. Each session comprised 8 trials. Sessions with fewer than 4 completed trials were excluded, yielding 271 analyzable sessions across 30 participants.

Task structure was held constant across sessions to preserve comparable demands, including the arena, number of chests, number of object-containing chests, distractor phase, retrieval procedure, feedback, and scoring. Objects were not repeated within a session. Across sessions, individual objects could recur, but object-location associations were newly generated because chest locations were randomized for each task instance. Thus, repeated sessions exposed participants to the same task format but not to the same object-location pairings.

### Study Procedures

The asm-EMA session was delivered on a guided-access tablet (iPad Pro 11-inch, 2nd generation). Sessions were scheduled automatically by a custom server between 9:00 AM and 9:00 PM. Following study initiation, the first asm-EMA session was delivered after a random delay of 20–30 minutes, and subsequent asm-EMA sessions were delivered at randomized intervals of 90–150 minutes (i.e. 2 hours +/- 30 minutes). Each session began with an audiovisual startle cue consisting of a 50-ms, 95-dB burst of white noise and a flashing tablet screen. The alert remained active while participants completed the first three required items, (current activity, VAS Pain, and VAS Anxiety) and stopped immediately after the VAS Anxiety response, before administration of the optional STAI-6 items and Treasure Hunt memory task. Responses were stored on a password-protected, HIPAA-compliant server accessible only to authorized study investigators.

### Baseline and Follow-up Questionnaires

At the beginning and end of the inpatient hospitalization period, subjects underwent a baseline evaluation of psychosocial wellbeing and environmental spatial ability by completing the Beck Anxiety Inventory (BAI), Beck Depression Inventory-II (BDI-II), Quality of Life in Epilepsy-31 (QOLIE-31), Spielberger State-Trait Anxiety Inventory (STAI), and Santa Barbara Sense of Direction scale (SBSOD). To account for possible fluctuations in trait-level psychosocial wellbeing and spatial ability survey responses due to impending surgery or hospital stay, the baseline battery was administered twice: once at the beginning of hospitalization, typically on day following enrollment, and again at the end of hospitalization. Subjects were initially instructed to complete the second battery remotely within 2 weeks of discharge from the hospital; however, due to inadequate participation rates, subsequent patients were administered the second battery on the day prior to discharge.

The full-form Spielberger State-Trait Anxiety Inventory (STAI) is a 40-item assessment consisting of two 20-item questionnaires, one of which measures state anxiety (Form Y-1) and the other, trait anxiety (Form Y-2). State anxiety captures a person’s immediate level of anxiety within a particular context or moment in time, whereas trait anxiety reflects a person’s general predisposition to anxiety and can be influenced by both prior and potential future experiences with anxiety^42^. Each form of the STAI includes 20 statements that describe an individual, and subjects are asked to indicate the extent to which each statement applies to them, ranging from 1 (almost never) to 4 (almost always). Select items are reverse-scored, and the final score for each form is calculated by summing the scores across all 20 questions, using the corrected score for reverse-scored items. Scores for each form of the STAI can range from 20 to 80, with higher scores indicating a greater level of anxiety^43^.

The Beck Anxiety Inventory (BAI) is a 21-item self-report questionnaire that assesses the severity of a subject’s anxiety. Unlike the STAI, this survey was specifically designed to discriminate between anxiety and depression, and to serve as a reliable measure of anxiety in clinical psychiatric populations^44^. Each question describes a common symptom of anxiety, and subjects are asked to rate how much they have been bothered by each symptom during the past week. Responses range from 0 (“Not at all”) to 3 (“Severely – I could barely stand it”). Individual item scores are then summed to yield a total score ranging from 0-63, with 0-7 suggesting minimal anxiety, 8-15 mild anxiety, 16-25 moderate anxiety, and 26-63 severe anxiety^45^.

The Beck Depression Inventory-II (BDI-II) is a 21-item self-report inventory used to assess the severity of a patient’s depression^46^. Each item begins with a topical header that indicates the general area of focus for the statements to follow. Subjects are then presented with four statements listed in increasing severity and are asked to choose the one statement that best represents the way they have been feeling over the past two weeks. Individual items are scored from 0-3 and summed for a total score ranging from 0 to 63, with higher scores indicating greater severity of depression. Scores from 0-13 reflect minimal depression, 14-19 mild depression, 20-28 moderate depression, and 29-63 severe depression^47^.

The Quality of Life in Epilepsy Inventory (QOLIE-31) is a 31-item questionnaire that measures the health-related quality of life in adults with epilepsy. The survey includes 1 question that assesses overall health and 30 questions which each fall into one of seven categories or multi-item “scales”: seizure worry, overall quality of life, emotional well-being, energy/fatigue, cognitive, medication effects, and social function. Each multi-item scale produces a scale score that reflects the average response across all items within that category. The overall QOLIE-31 score is obtained using a weighted average of the multi-item scale scores and can range from 0-100, with a higher overall score indicating a higher quality of life^48^.

The Santa Barbara Sense of Direction Scale is a 15-item self-report questionnaire that measures a subject’s perceived spatial orientation at the environmental level^49^. Each item provides a statement about a subject’s spatial and navigational abilities, preferences, or experiences, and subjects are asked to indicate their level of agreement with the statement on a scale of 1 (strongly agree) to 7 (strongly disagree). Seven of 15 items are reverse-scored, and the final score is obtained by averaging the score across all 15 items, using the corrected score for reverse-scored items. A higher final score indicates a stronger perceived sense of direction.

### Data Quality and Exclusion Criteria

Thirty-three participants were recruited for the study. Three participants had no usable EMA data and were excluded from analysis, yielding a final analytic sample of 30 participants. Sessions were excluded if the VAS pain or VAS anxiety ratings were not completed, or if fewer than 4 Treasure Hunt trials were completed. For the STAI-6, two additional response quality checks were applied. First, sessions were excluded if a participant gave identical responses to all six items, either all 1s or all 4s, as this pattern indicates uniform responding without regard to question content. Second, sessions were excluded if the time elapsed between consecutive STAI-6 item responses was less than 1 second throughout, suggesting the questions were not read. In total, 7 sessions were excluded based on these criteria, and the remaining 264 sessions were used in all analyses involving STAI-6.

### Descriptive distributions, variance decomposition, and clinical validity

First, we characterized the empirical distribution of each asm-EMA measure across completed sessions and quantified the relative contributions of between-subject and within-subject variation using intraclass correlation coefficients (ICCs) estimated from random-intercept linear mixed-effects models. This descriptive step was used to summarize the extent to which each asm-EMA measure reflected stable individual differences versus session-to-session fluctuation.

Linear mixed-effects models were represented using the general form:

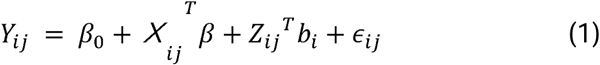

where *Y_ij_* is the outcome for participant *i* at session *j,X_ij_* contains the fixed-effect predictors, and *Z_ij_^T^*, *b_i_* represents the participant-specific random effects. All models included a participant-specific random intercept, with random slopes included where specified below. The specific fixed- and random-effect structures for each analysis are provided using model-formula notation:

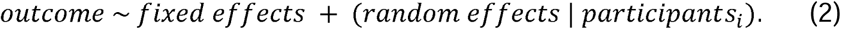

For each asm-EMA measure, the ICC model was fitted by restricted maximum likelihood and specified as *raw measure∼*1 + (1 | *participant*) The ICC was computed as:

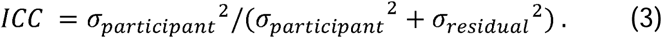

Second, to evaluate clinical validity, we computed Spearman correlations between subject-level asm-EMA summaries and pre- and post-monitoring neuropsychiatric trait assessments. For each asm-EMA measure, we computed two subject-level summaries from each participant’s repeated sessions. The subject-level mean reflected each participant’s average level across the monitoring period, and the subject-level standard deviation across sessions reflected each participant’s session-to-session variability. These analyses tested whether average asm-EMA responses and asm-EMA variability were associated with established clinical assessments collected before and after the study.

For analyses focused on within-person variation, each asm-EMA observation was transformed into a person-relative value: *Y*_i,j_* = (*Y_ij_* − <u>*Y_ij_*</u>)/ where (*<u>Y</u>*)*s_Y,centered_*, is the participant-specific mean for that measure and *s_Y,centered=_SD({Y_ij_-<u>Y</u>}_ij_)* is the standard deviation of all person-mean-centered observations for that measure pooled across participants. This transformation removed stable between-person differences in average level while preserving differences across participants in how much they fluctuated over time. Unless otherwise stated, within-subject correlations, within-session mixed-effects models, and temporal dynamics analyses used these person-relative values.

### Between-subject correlation analyses

Third, to characterize associations among the asm-EMA measures at the between-subject level, we computed Spearman correlations using subject-level means. These analyses tested whether participants with higher average anxiety also exhibited different average memory performance. Because one mean per participant and measure was entered into the analysis, these correlations reflected stable between-subject differences rather than session-to-session covariation.

### Within-session associations and linear mixed-effects models

To characterize associations among the asm-EMA measures at the within-subject level, we computed Spearman correlations using person-relative session-level observations. These analyses tested whether session-specific deviations from a participant’s typical anxiety level covaried with corresponding deviations in memory performance within the same asm-EMA session.

To formally test within-subject anxiety–memory relationships while accounting for repeated observations nested within participants, we fitted linear mixed-effects models following the general framework defined above. Memory accuracy and response time were modeled separately, with each affective measure entered as a fixed-effect predictor and participant included as a random intercept:

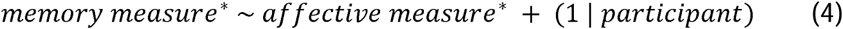

Here, the outcome was person-relative Memory Accuracy or Memory RT, and the predictor was person-relative VAS Pain, VAS Anxiety, or STAI-6 Anxiety. To test whether any affective measure explained unique variance beyond the others, we also fitted multivariable models including all three affective predictors:

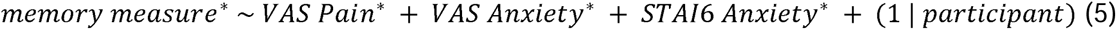

### Temporal dynamic analysis

Last, to test whether asm-EMA measures showed short-term temporal structure, we examined changes across repeated assessments. Only within-day lagged pairs were retained; cross-day pairs and same-day pairs separated by more than five hours were excluded. Within-person Spearman autocorrelations were computed for each asm-EMA measure at lags 1, 2, and 3, testing whether person-relative values of the same measure were correlated across adjacent and later sessions. We also computed within-person Spearman cross-correlations for all ordered pairs of asm-EMA measures to quantify lagged coupling between different measures. Both autocorrelations and cross-correlations were computed using the person-relative values defined above.

For all temporal mixed-effects analyses described below, model performance was summarized using R², and statistical significance was evaluated using 2,000 within-subject permutations. The specific predictor or predictors shuffled to generate each permutation null are described for each analysis below.

Before modeling session-to-session predictability, we separately quantified change across hospitalization. For each measure, we fitted a linear mixed-effects model with participant-specific random intercepts and hospital-day slopes:

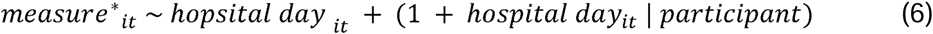

Statistical significance was evaluated against a within-subject permutation null generated by shuffling hospital-day labels within each participant while retaining the same model specification. A significant R² indicated that the observed association with hospital day was stronger than expected when the within-participant ordering of hospital days was disrupted.

Prior to fitting mixed autoregressive models, we applied per-subject linear detrending to remove these gradual hospital-day trends. For each participant separately, we fitted:

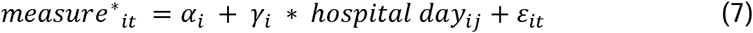

and retained the residuals *measure_it_* = *measure_it_*\* - (*α_i_* + *γ_i_* * *hospital day_it_*) as the detrended series. We then fitted mixed autoregressive models on the detrended scores separately for each asm-EMA measure:

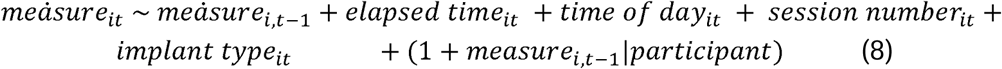

Models were fitted by maximum likelihood and summarized using R². The permutation null was generated by shuffling the same-measure lag-one predictor within each participant while retaining the outcome, covariates, and model specification unchanged. A significant *R*^2^ indicated that the lag-one term improved prediction beyond the covariates relative to a within-participant shuffled lag-one null, while *β_lag1_* quantified the direction and magnitude of short-term persistence.

To test whether one affective measure predicted later values of another beyond the outcome’s own short-term persistence, we extended the autoregressive model for the three affective outcomes (VAS Pain, VAS Anxiety, STAI-6) by adding the two lag-one detrended scores from the remaining affective measures as cross-predictors:

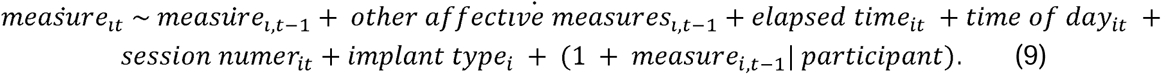

Models were evaluated using the same within-subject permutation framework. Here, the same-measure lag-one predictor was retained in its original temporal alignment, and only the two cross-predictors were shuffled within each participant. A significant 7 indicated that the lagged values of other affective measures improved prediction beyond the outcome’s own lag-one term and the covariates, while their coefficients quantified the direction and magnitude of these cross-measure associations.

## Declaration Statements

### Data Availability

The datasets generated and/or analyzed during the current study are not publicly available due to pending formatting and privacy requirements for an NIH sponsored data archive but are available from the corresponding author on reasonable request.

### Code Availability

The code generated and/or analyzed during the current study is publicly available at https://github.com/claire98han/EMAanalysis.

### Abbreviations

EMA: ecological momentary assessment
asm-EMA: anxiety-spatial-memory EMA
VAS: Visual Analog Scale
STAI: Spielberger State-Trait Anxiety Inventory
STAI-6: six-item STAI
BAI: Beck Anxiety Inventory
BDI-II: Beck Depression Inventory-II
QOLIE-31: Quality of Life in Epilepsy-31
SBSOD: Santa Barbara Sense of Direction scale
EMU: Epilepsy Monitoring Unit
EEG: electroencephalography
SEEG: stereoelectroencephalography
ICC: intraclass correlation coefficient
RT: response time.

### Author Contributions

BEY, JJ, HZ, KCZ, YL, CZH, LDK, RPA, LMW, MH and CAS designed the study. KCZ, HZ, AGV, SS, AC and JC collected the data. CZH and LMW did data pre-processing. CZH, KCZ and LMW analyzed the data. CZH, KCZ and LMW drafted and all authors contributed to the manuscript. All authors have read and approved the manuscript.

## Acknowledgements

We would like to thank all patients for their participation, clinical faculty and staff from Columbia University Irving Medical Center and NewYork-Presbyterian Hospital for support with subject recruitment and participation.

