## Supplemental material for "Investigating the Dynamic Relationship Between Anxiety and Spatial Memory Using Autonomous Ecological Momentary Assessment"

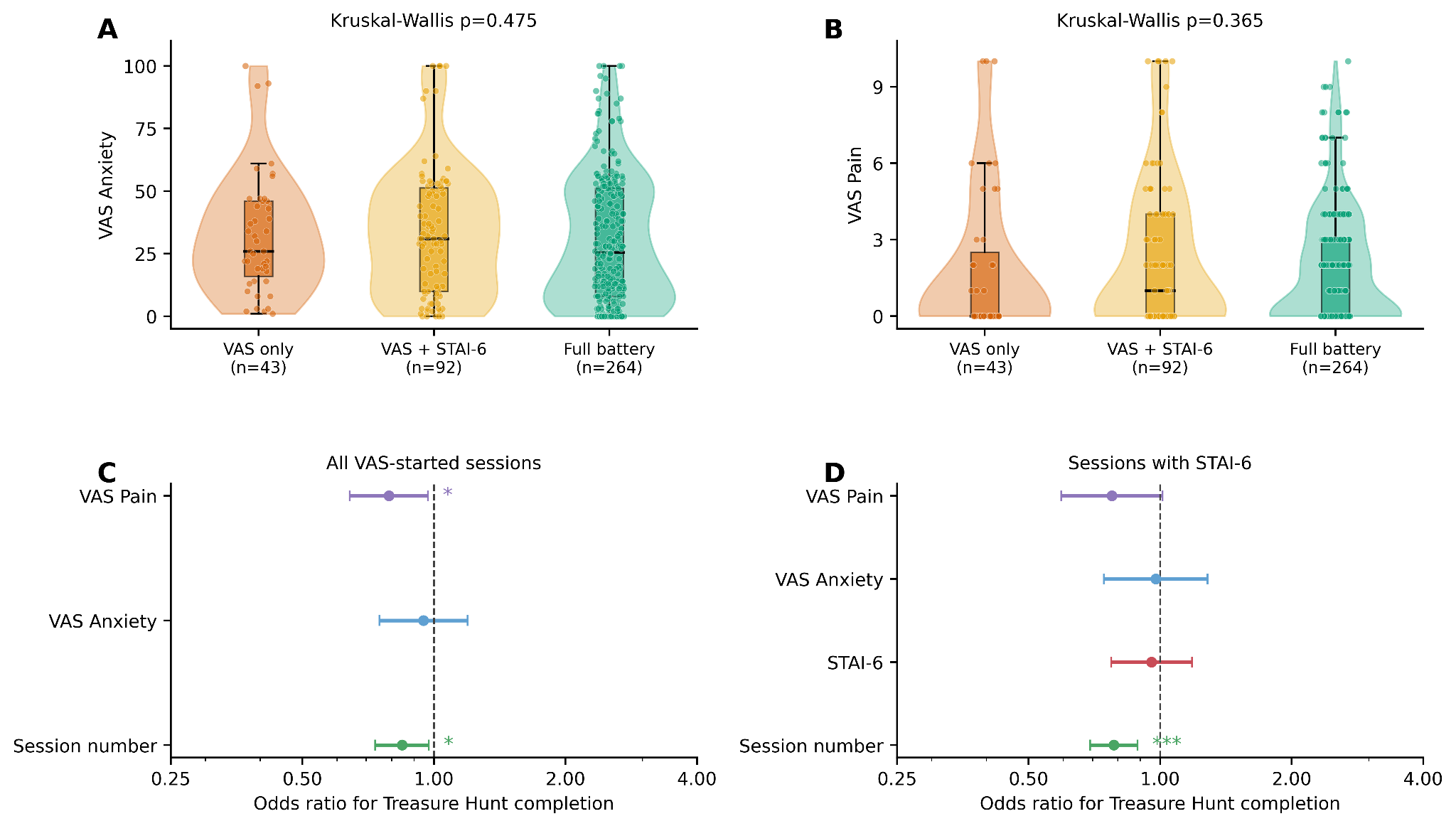

**Figure S1. Session-level analyses of EMA completion and missing Treasure Hunt data.** (A, B) VAS Anxiety and VAS Pain ratings are shown across session completion categories, including sessions in which participants completed only the initial VAS ratings, completed VAS + STAI-6, or completed the full battery including the Treasure Hunt memory task. Violin plots show score distributions, overlaid with box plots and jittered session-level data points. Kruskal-Wallis tests showed no significant differences across completion categories for either VAS Anxiety or VAS Pain. (C, D) Session-level sensitivity analyses testing whether momentary affective state predicted usable Treasure Hunt completion. Odds ratios are shown from logistic models accounting for repeated sessions within participants. Panel C includes all sessions with valid VAS ratings, and Panel D includes the subset of sessions with STAI-6 data. VAS Anxiety and STAI-6 did not predict Treasure Hunt completion, whereas higher VAS Pain and later session number were associated with lower odds of completion. Error bars indicate 95% confidence intervals. Significance markers indicate p < 0.05 and p < 0.001.

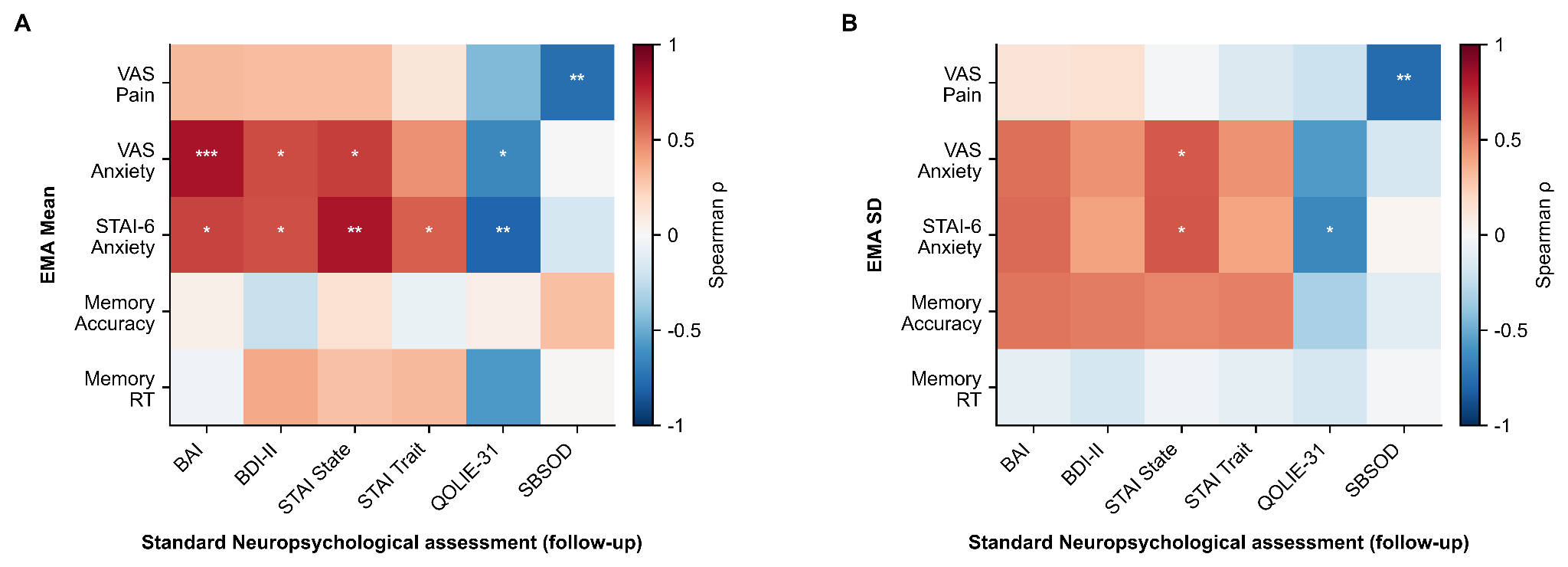
 **Figure S2. Correspondence between asm-EMA measures and post-EMA clinical assessments.** Spearman correlation matrices between post-EMA clinical assessment scores (x-axis: BAI, BDI, STAI State, STAI Trait, QOLIE-31, SBSOD) and subject-level asm-EMA means (A) and asm-EMA within-person standard deviations (B) for all five measures (VAS Pain, VAS Anxiety, STAI-6, Memory Accuracy, Memory RT). Color encodes the Spearman ρ value; asterisks indicate significance (* p < .05, ** p < .01, *** p < .001). These analyses parallel the pre-EMA assessment correlations reported in Figure 4 and should be interpreted cautiously given the reduced sample size at post-EMA period assessment.

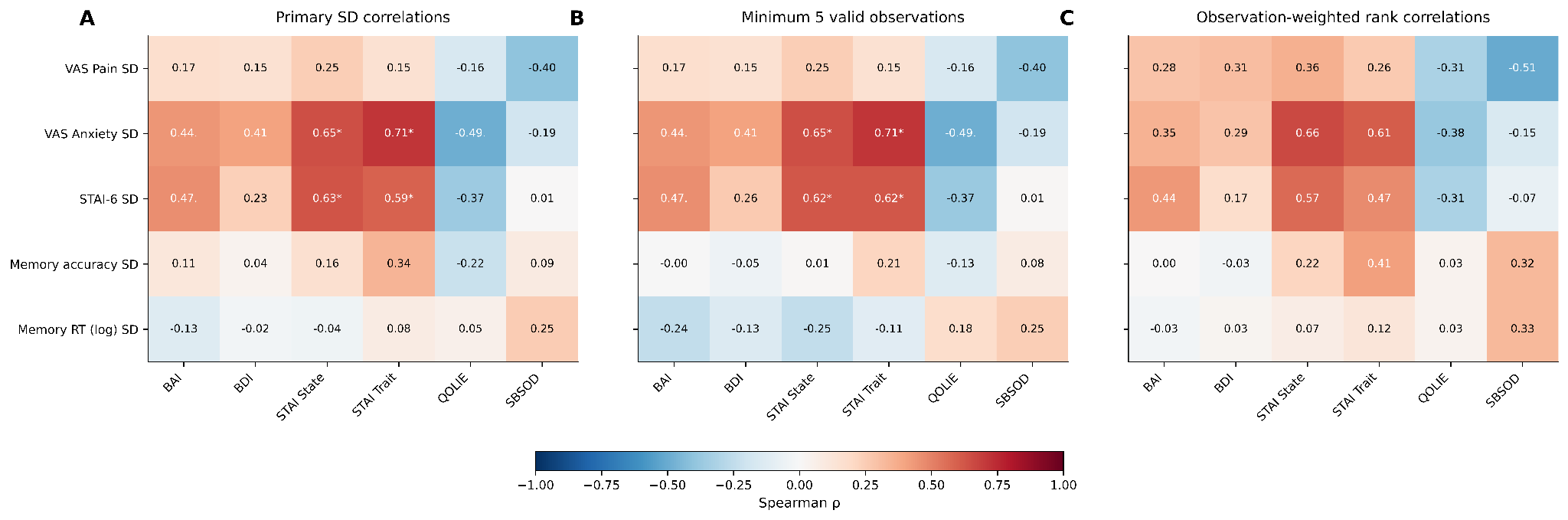
 **Figure S3. Sensitivity analyses for clinical validity correlations involving EMA variability.** (A) Primary Spearman correlations between participant-level EMA standard deviation estimates and baseline clinical measures. (B) Sensitivity analysis restricted to participants with at least five valid observations for the relevant EMA measure. (C) Observation-weighted rank correlations, weighting each participant by the number of observations contributing to the EMA standard deviation estimate. Cell values show Spearman rho and sample size. Significance markers indicate Benjamini-Hochberg FDR-adjusted q-values within each panel.

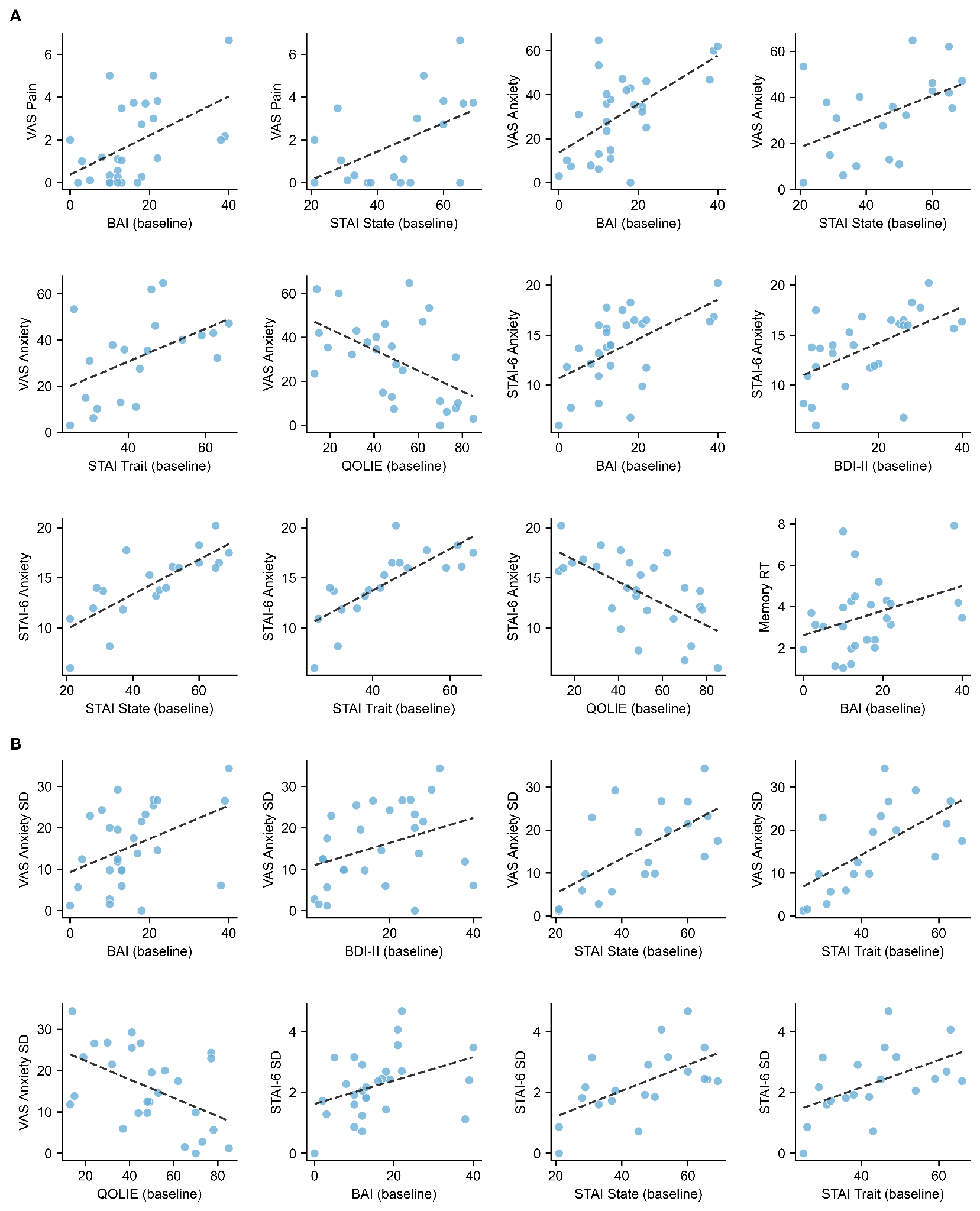

**Figure S4. Scatter plots of significant correlations between asm-EMA measures and standard neuropsychological assessments at baseline.** Each panel shows the Spearman correlation between one asm-EMA measure (y-axis) and one standard clinical instrument administered before the monitoring period (x-axis). Dashed lines show ordinary least-squares fits. (A) show correlations with subject-level asm-EMA means; (B) show correlations with within-person standard deviations (SD). Only statistically significant pairs (p < 0.05) from the corresponding heatmaps (Figure 4A–B) are displayed. BAI, Beck Anxiety Inventory; BDI-II, Beck Depression Inventory; STAI State and Trait, State-Trait Anxiety Inventory; QOLIE, Quality of Life in Epilepsy; SBSOD, Santa Barbara Sense of Direction scale.

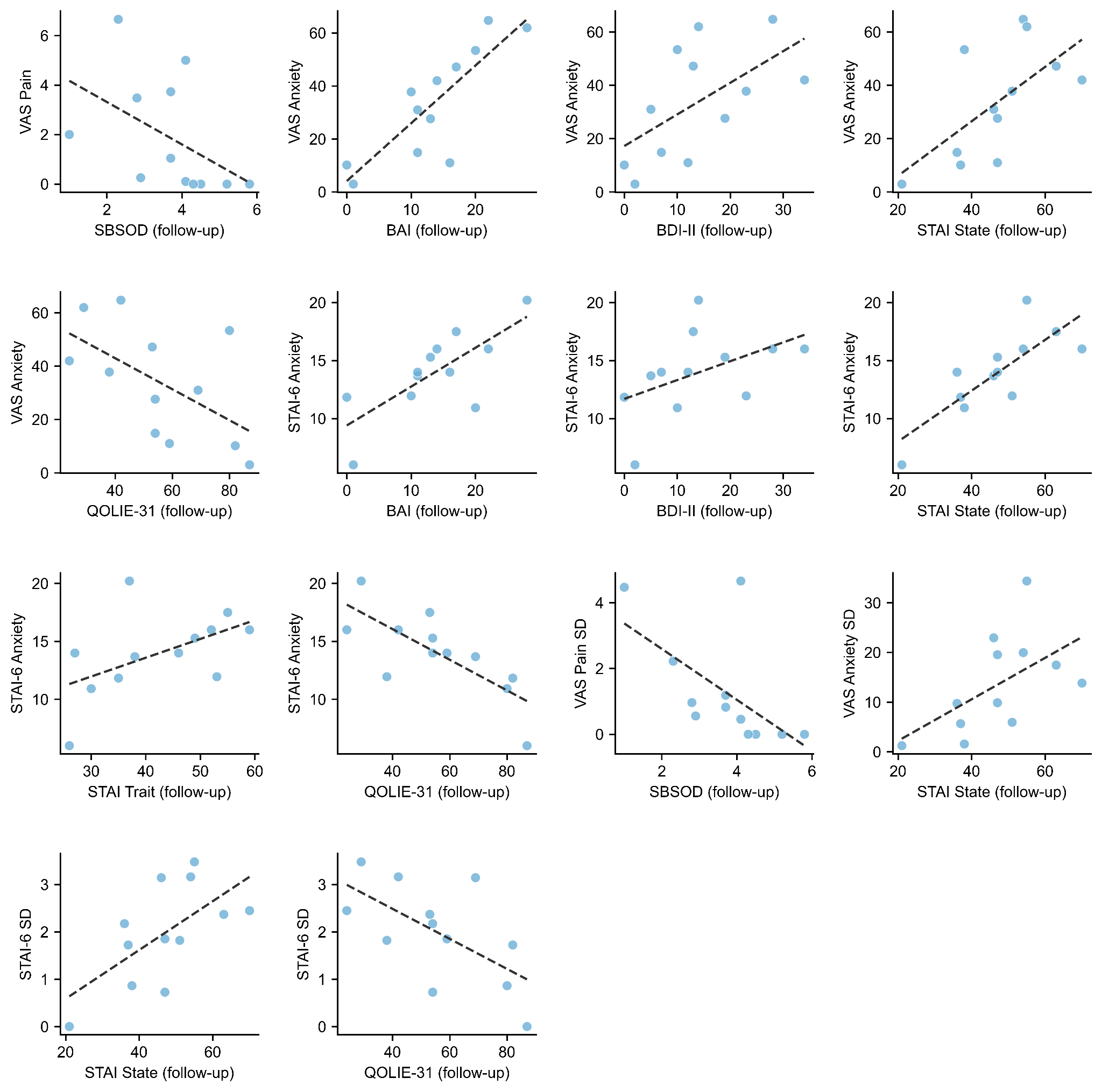

**Figure S5. Scatter plots of significant correlations between asm-EMA summary statistics and standard neuropsychological assessments at follow-up.** Each panel shows the Spearman correlation between one asm-EMA measure (y-axis) and one standard clinical instrument administered after the monitoring period (x-axis). Dashed lines show ordinary least-squares fits. Panels in the upper portion show correlations with subject-level asm-EMA means (VAS Pain, VAS Anxiety, STAI-6 Anxiety); panels in the lower portion show correlations with within-person standard deviations (VAS Pain SD, VAS Anxiety SD, STAI-6 SD). Only statistically significant pairs (p < 0.05) from the corresponding follow-up heatmaps are displayed. Note that sample size is reduced relative to baseline analyses owing to missing follow-up assessments in a subset of participants.

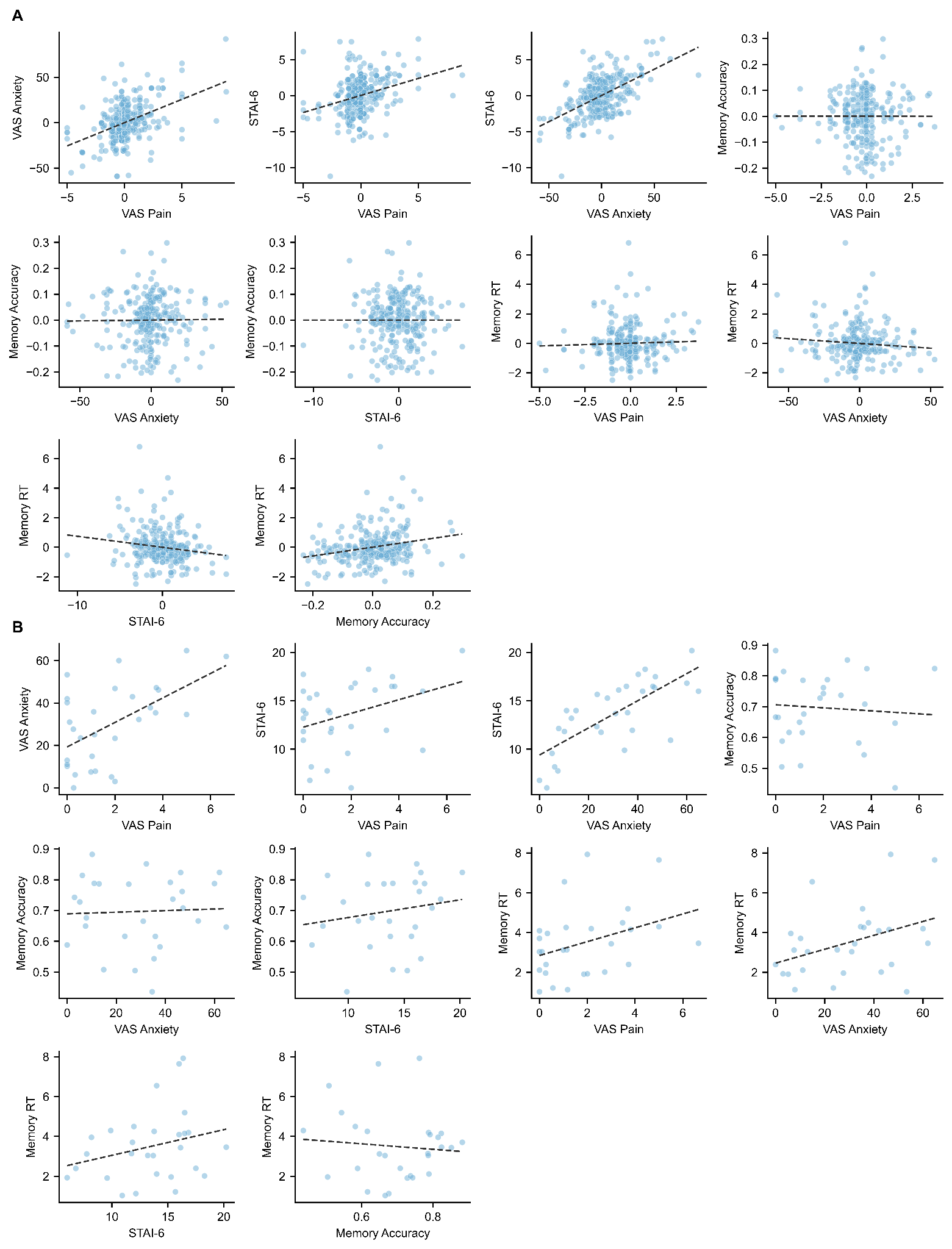

**Figure S6. Pairwise scatter plots of all asm-EMA measures at the within-person and between-person levels.** Each panel shows the relationship between two asm-EMA measures with a least-squares regression line overlaid. Panel A shows within-person associations computed on person-mean centered session-level data, where each data point represents a single asm-EMA session expressed as a deviation from that participant's own mean. Panel B shows between-person associations computed on subject-level means, where each data point represents one participant's average across all their sessions. The 10 panels in each section correspond to all pairwise combinations of the five asm-EMA measures (VAS Pain, VAS Anxiety, STAI-6, Memory Accuracy, Memory RT), covering the lower triangle of the full correlation matrix. These scatter plots provide individual-level visualization of the Spearman correlations summarized in the heatmap in Figure 5A.

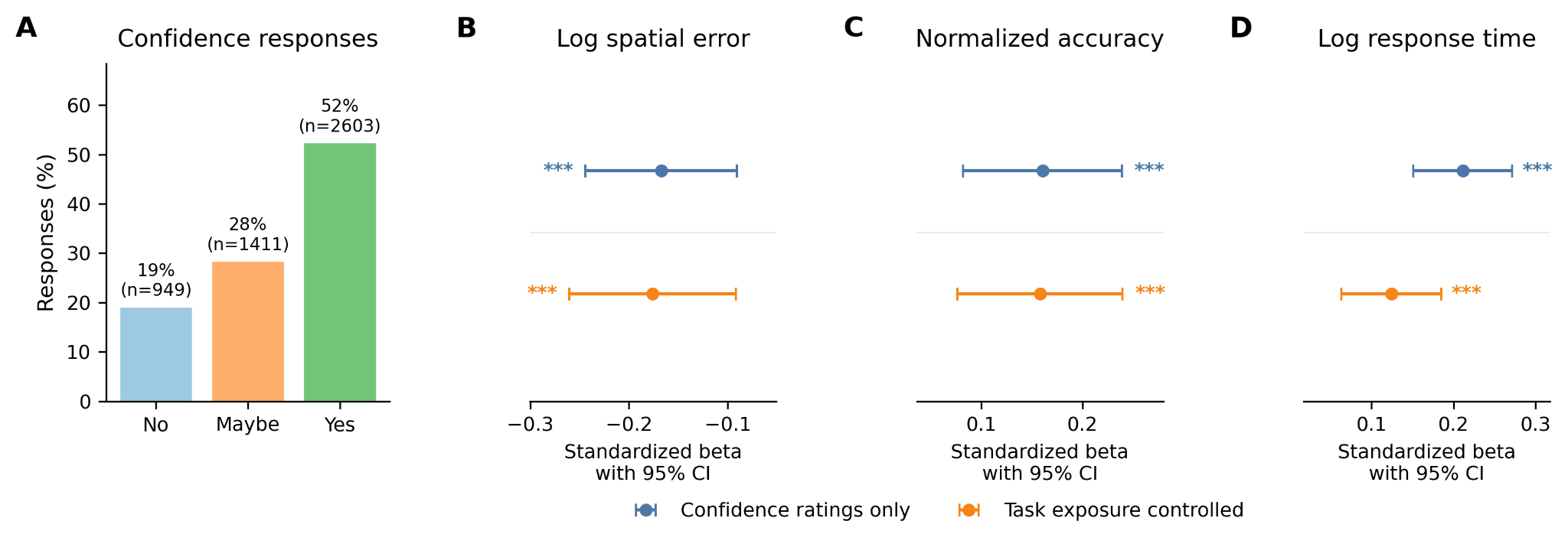

**Figure S7. Confidence ratings track Treasure Hunt task behavior.** (A) Distribution of confidence responses across Treasure Hunt retrieval trials. Bars show the percentage of responses rated “No,” “Maybe,” or “Yes,” with counts shown above each bar. (B–D) Mixed-effects models testing associations between within-person confidence ratings and task behavior, including log spatial error (B), normalized memory accuracy (C), and log response time (D). Blue points show models including confidence ratings only, and orange points show models additionally controlling for prior task exposure. Points indicate standardized beta coefficients, and error bars indicate 95% confidence intervals. Significance markers indicate p < 0.05, p < 0.01, and p < 0.001.

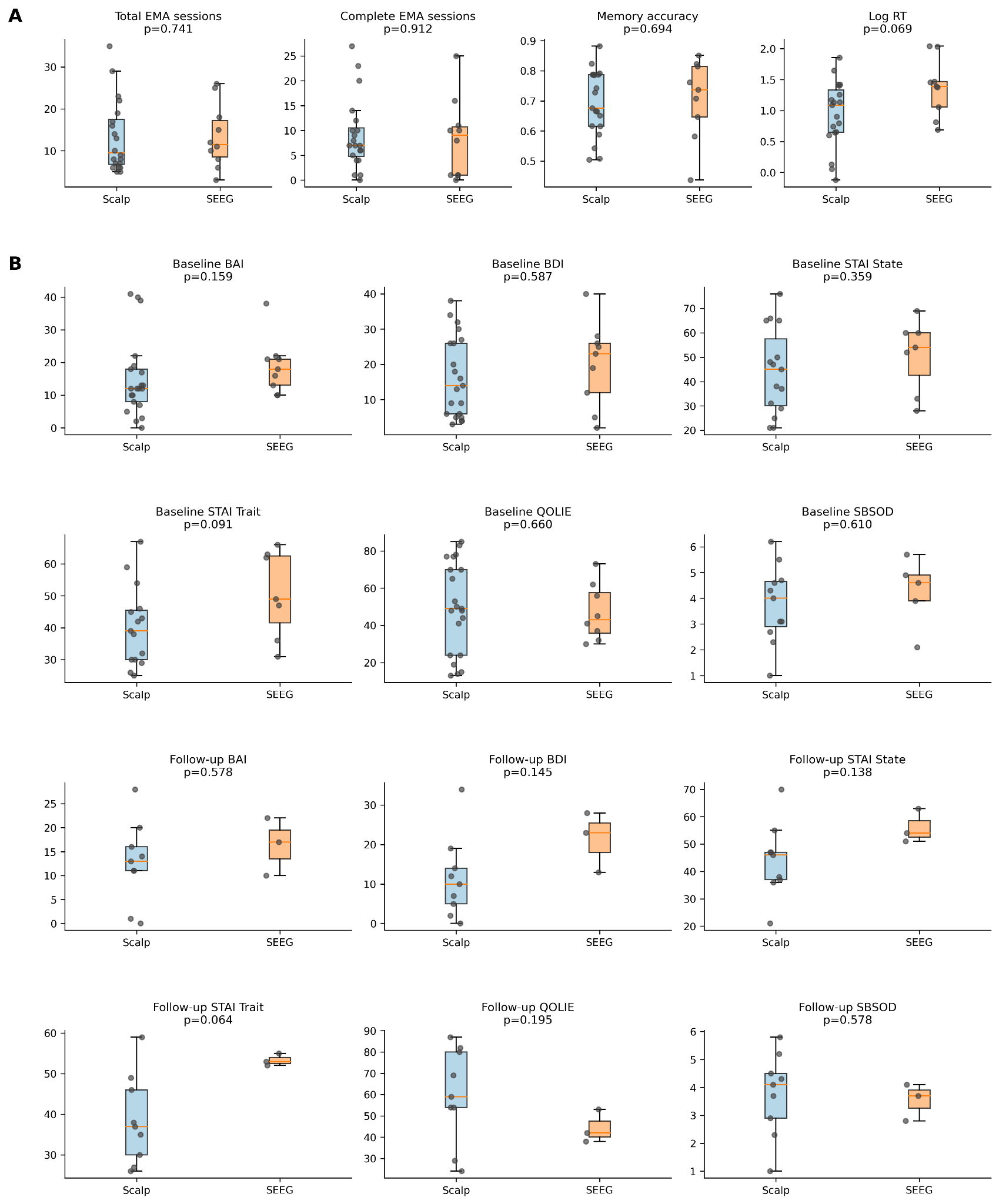

**Figure S8. Sensitivity analyses comparing SEEG and scalp EEG participants.** (A) Group comparisons for EMA participation and task performance, including total EMA sessions, complete EMA sessions, memory accuracy, and log-transformed memory response time. (B) Group comparisons for baseline and follow-up neuropsychological measures. Each point represents one participant, boxplots summarize the group distributions, and p-values reflect SEEG versus scalp EEG group comparisons.

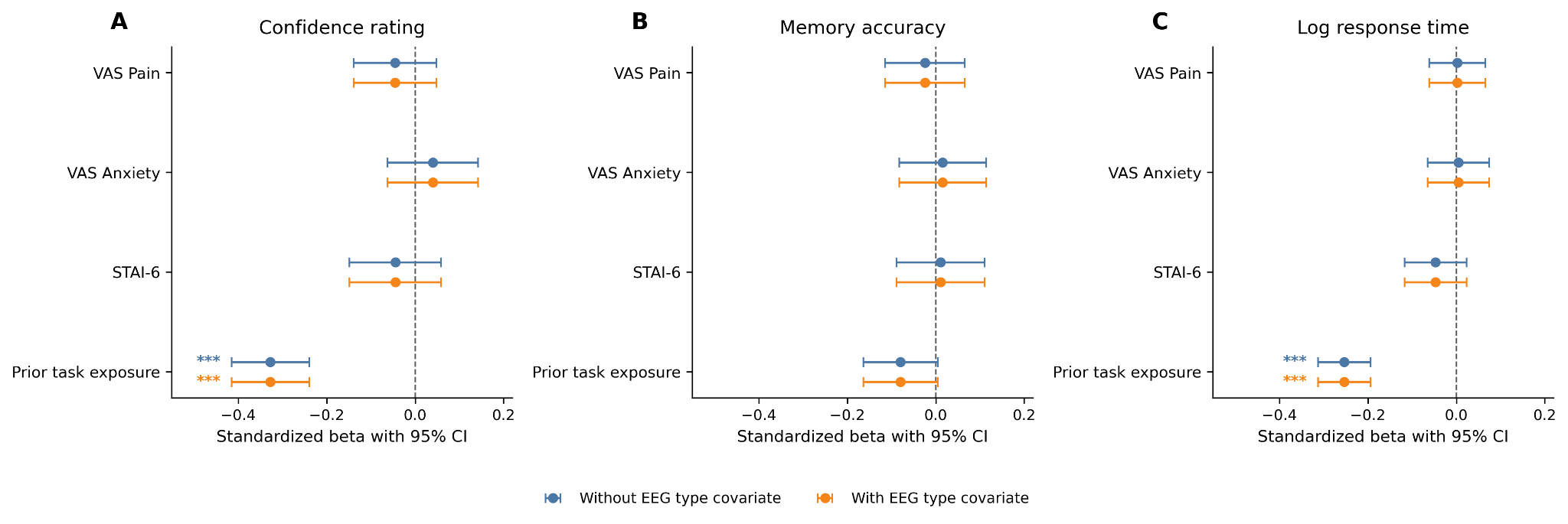

**Figure S9. EEG type covariate sensitivity analysis for the primary mixed-effects models.** Models were fit separately for confidence rating (A), memory accuracy (B), and log-transformed response time (C). Points show standardized beta coefficients and error bars indicate 95% confidence intervals. Blue estimates show models without EEG type as a covariate, and orange estimates show models additionally adjusting for EEG type. Including EEG type did not materially change the affective predictor estimates or the prior task exposure effect. Significance markers indicate p < 0.05, p < 0.01, and p < 0.001.

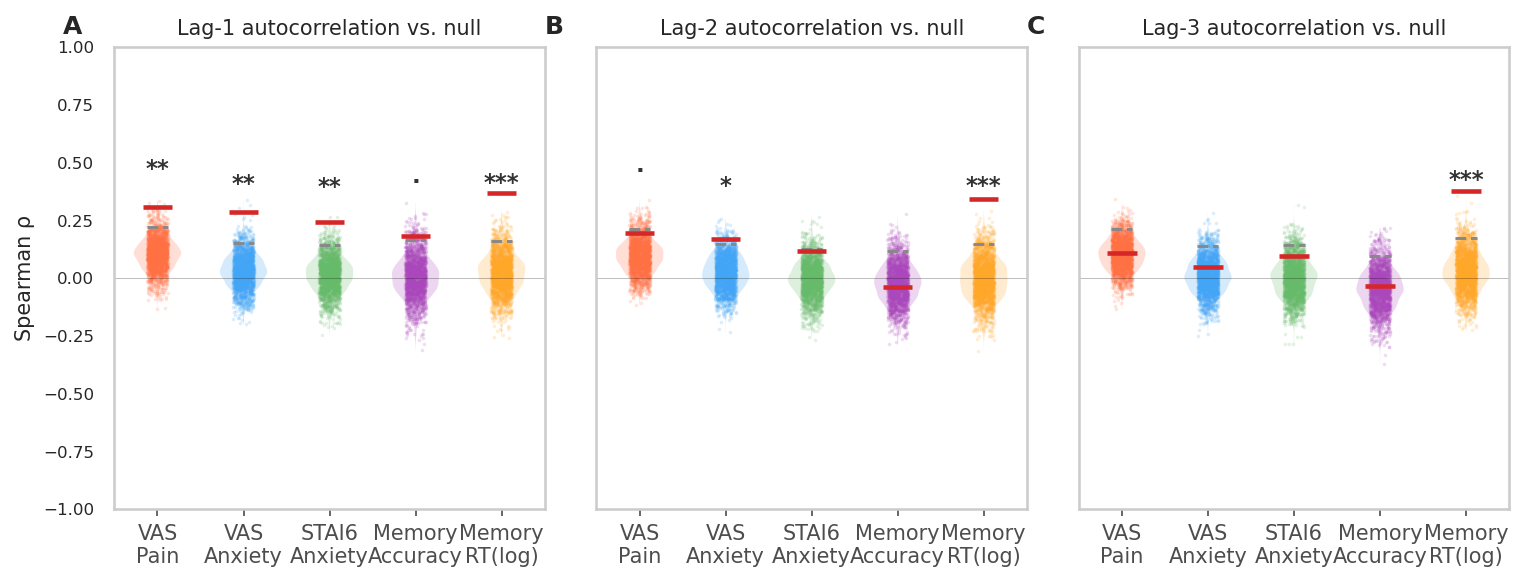

**Figure S10. Null distributions for lag-1, lag-2, and lag-3 autocorrelations.** Each shaded area shows the permutation null distribution (N = 2000 within-subject shuffles) for pooled Spearman ρ for each measure. Red dashes indicate observed ρ; dashed grey lines indicate the 95th null percentile. Asterisks above violins indicate permutation significance (* p < .05, ** p < .01, *** p < .001). All five measures are shown; the y-axis is shared across lags.

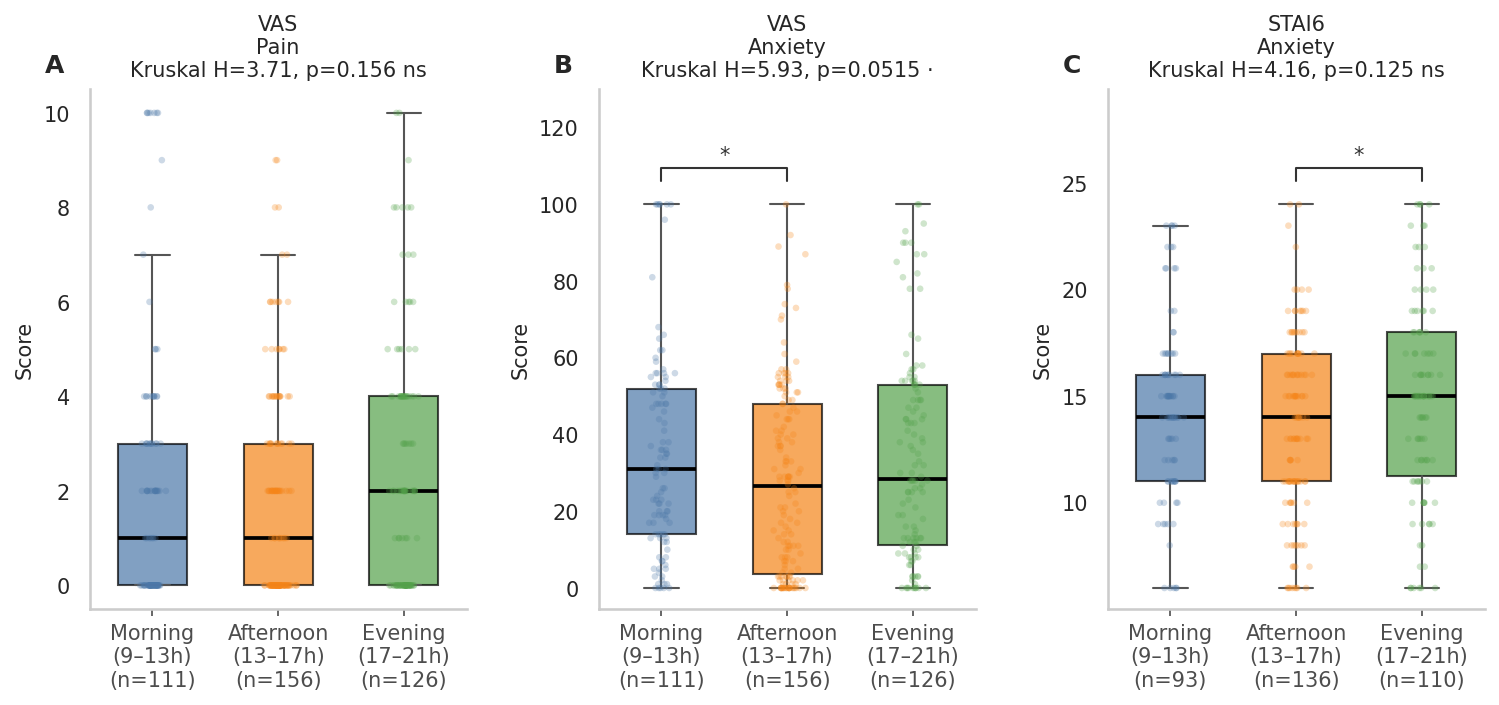

**Figure S11. Session-level** asm-**EMA scores by time of day (raw scores).** Violins show the distribution of session scores in morning (9–13h), afternoon (13–17h), and evening (17–21h) windows. Medians are shown as horizontal bars; individual sessions are jittered. Brackets indicate pairwise Mann-Whitney comparisons significant at uncorrected p < .05 (* p < .05). Overall Kruskal-Wallis H and p are reported in each panel title.

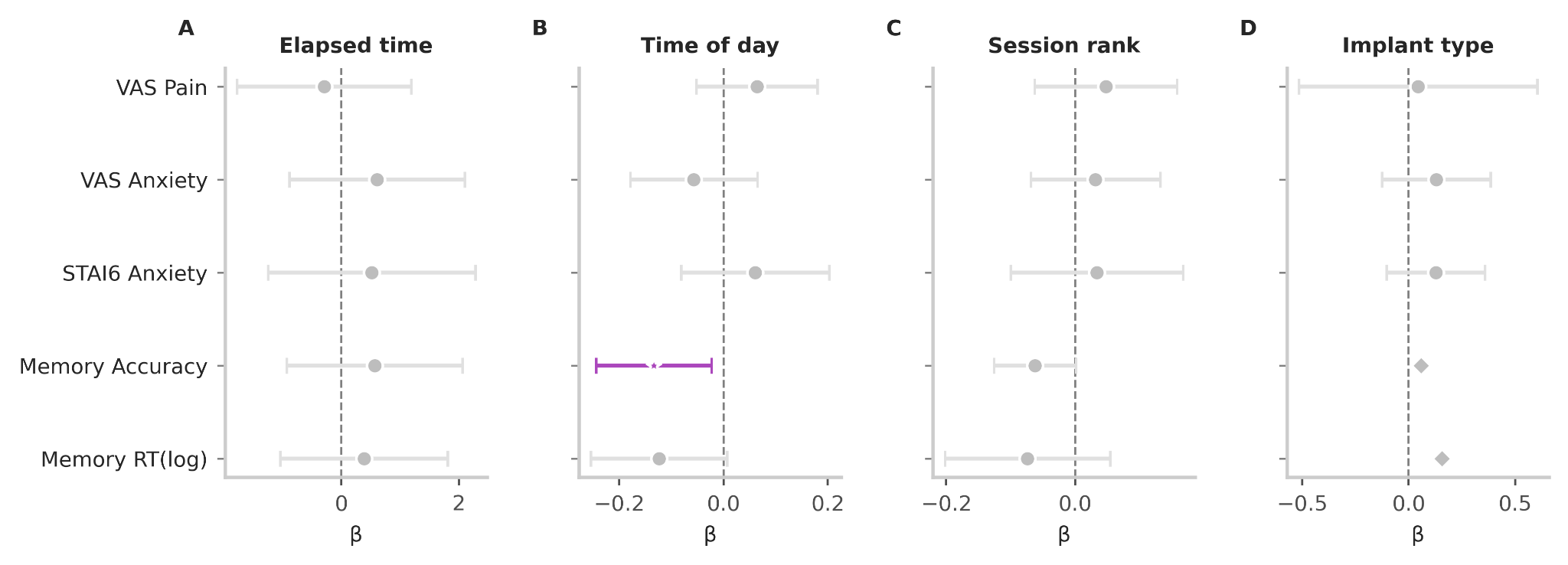

**Figure S12**: **β coefficients (± 95% CI) for elapsed time between sessions (A), time of day (B), session rank (C), and implant type (D) from mixed-effects lag-1 AR models fitted for each of the five measures.** "Outcome measures were person-mean-centred and scaled by the pooled within-person standard deviation prior to modelling (see Methods); Grey markers indicate non-significant effects (p ≥ .05); colored markers indicate p < .05. Among all covariates, time of day showed the most consistent pattern: both memory measures exhibited significant negative associations (Accuracy: β = −0.165, p = .018; RT: β = −0.133, p = .013), indicating that accuracy declined and responses were faster later in the day, whereas affective measures were unaffected by time of day.

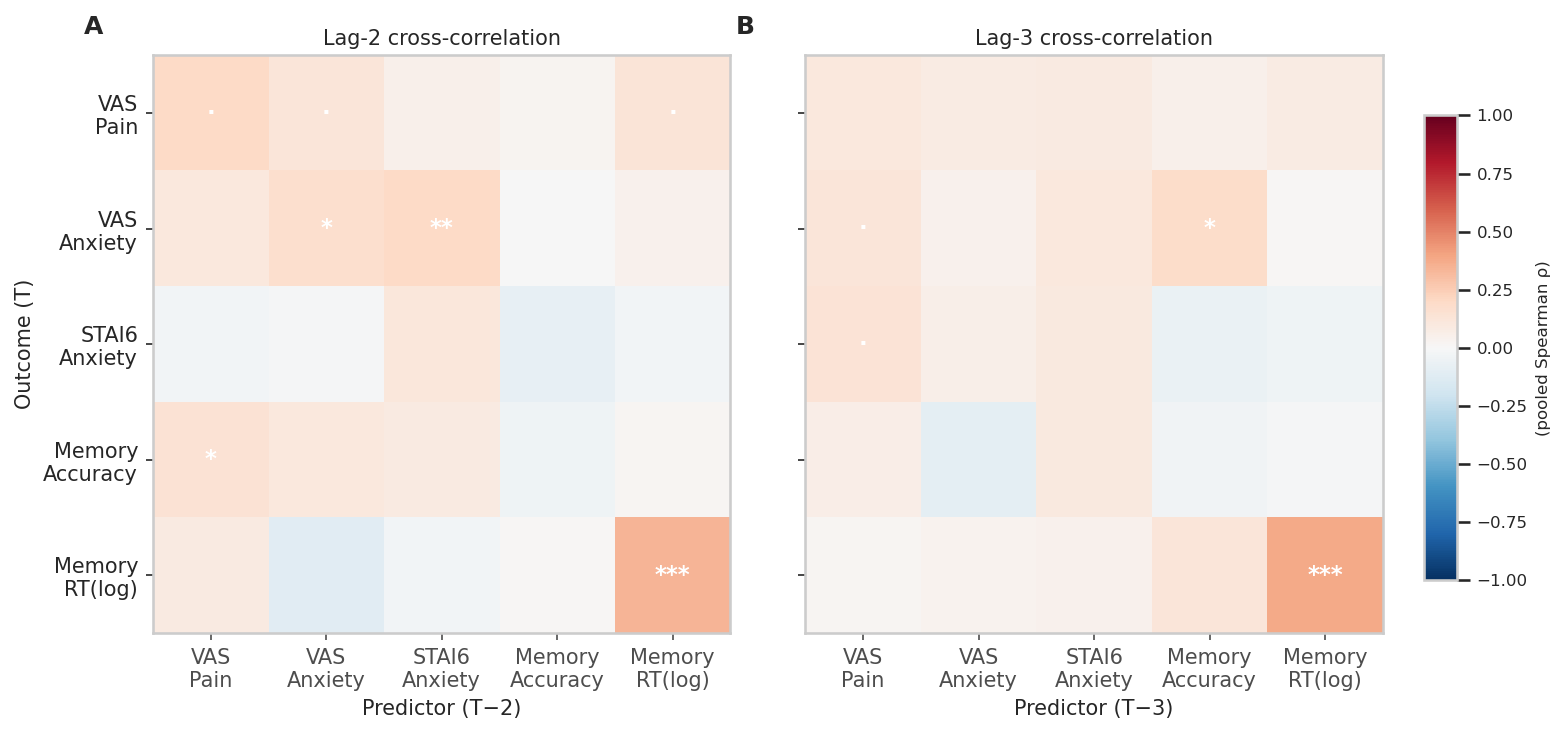

**Figure S13**: **Lag-2 and lag-3 cross-correlation matrices (pooled Spearman ρ).** Same format as Figure 6B. Rows = outcome at T, columns = predictor at T−2 (left) or T−3 (right). Asterisks denote permutation significance (· p < .10, * p < .05, ** p < .01, *** p < .001). Y-axis labels shown for lag-2 only; lag-3 shares the same row ordering.

**Lag-2 Autoregression Permutation Testing results**

| asm-EMA measures | Observed R^2^ | Null 95 pct R^2^ | P (shuffle) |
| --- | --- | --- | --- |
| VAS Pain | 0.477 | 0.487 | 0.171 |
| VAS Anxiety | 0.341 | 0.376 | 0.577 |
| STAI6 Anxiety | 0.296 | 0.255 | 0.009 |
| Memory Accuracy | 0.196 | 0.233 | 0.202 |
| Memory RT (log) | 0.341 | 0.369 | 0.126 |

**Lag-3 Autoregression Permutation Testing results**

| asm-EMA measures | Observed R^2^ | Null 95 pct R^2^ | P (shuffle) |
| --- | --- | --- | --- |
| VAS Pain | 0.473 | 0.514 | 0.858 |
| VAS Anxiety | 0.346 | 0.367 | 0.352 |
| STAI6 Anxiety | 0.427 | 0.477 | 0.932 |
| Memory Accuracy | 0.113 | 0.205 | 0.345 |
| Memory RT (log) | 0.256 | 0.364 | 0.442 |

**Table S1. Incremental contribution of lag-2 (with lag-1 retained) and lag-3 predictors.** R² values are computed from the full multi-lag model (1 − RMSE²/Var(y)). Permutation p-values test whether the additional lag coefficient — lag-2 or lag-3, shuffled independently while preserving all other lags — significantly improves prediction beyond the preceding model. With the exception of STAI-6, which showed a significant lag-2 contribution (p = .009), no additional lag significantly improved prediction for any other measure (all p > .12). The lag-3 model did not improve prediction for any measure (all p > .34).

| **asm-EMA measures** | Day Trend R**^2^** | Day trend p (perm) | detrend-AR R^2^ | detrend-AR p (perm) | detrend-AR lag-1 β | 95% CI | detrend-AR lag-1 p | N pairs | N subject |
| --- | --- | --- | --- | --- | --- | --- | --- | --- | --- |
| VAS Pain | 0.041 | 0.051 | 0.295 | 0.161 | +0.016 | [-0.135, +0.168] | 0.831 | 253 | 30 |
| VAS Anxiety | 0.050 | 0.014 | 0.324 | 0.000 | +0.182 | [+0.007, +0.357] | 0.042 | 253 | 30 |
| STAI-6 Anxiety | 0.147 | 0.000 | 0.192 | 0.026 | +0.062 | [-0.186, +0.309] | 0.626 | 200 | 29 |
| Memory Accuracy | 0.153 | 0.001 | 0.185 | 0.645 | -0.019 | [-0.124, +0.087] | 0.726 | 148 | 21 |
| Memory RT (log) | 0.365 | 0.000 | 0.219 | 0.059 | -0.088 | [-0.317, +0.141] | 0.449 | 148 | 21 |

**Table S2. Daily trend and autoregressive self-prediction at lag-1 (extended version of Table 2, additionally reporting sample sizes).** Day-Trend R² is the proportion of total within-person variance explained by a linear mixed-effects model with hospital day as the fixed predictor and participant-specific intercepts and slopes; p is the one-tailed permutation p-value (proportion of 2,000 null R² values ≥ observed R²). AR R² summarizes model performance for the lag-1 autoregressive model fitted to detrended scores (detrended lag-1 score, elapsed time, time of day, session rank, and implant type; random intercept and random lag-1 slope per participant), evaluated against a within-participant permutation null generated by shuffling the detrended same-measure lag-1 predictor. β lag-1 is the fixed-effect estimate for the lag-1 predictor from the mixed-effects model, with 95% confidence interval and analytic p-value. n pairs is the number of valid consecutive session pairs entering the AR model; n subj is the number of participants contributing at least one pair. Bold highlights statistical significance.
